# Chloroquine induces eryptosis in *P. falciparum-infected* red blood cells and the release of extracellular vesicles with a unique protein profile

**DOI:** 10.1101/2023.01.11.523595

**Authors:** Claudia Carrera-Bravo, Tianchi Zhou, Jing Wen Hang, Harshvardhan Modh, Fred Huang, Benoit Malleret, Matthias G Wacker, Jiong-Wei Wang, Laurent Renia, Kevin SW Tan

## Abstract

Malaria is a vector-borne parasitic disease that affects millions worldwide. In order to reach the objective, set by the World Health Organization to decrease the cases by 2030, antimalarial drugs with novel modes of action are required. Previously, a novel mechanism of action of chloroquine (CQ) was reported involving features of programmed cell death in the parasite, mainly characterized by calcium efflux from the digestive vacuole (DV) permeabilization. Increased intracellular calcium induces the suicidal death of erythrocytes also known as eryptosis. This study aimed to identify the hallmarks of eryptosis due to calcium redistribution and the downstream cellular effects during CQ treatment in iRBCs. *Plasmodium falciparum* 3D7 at mid-late trophozoites were used for the antimalarial drug treatment. Our results revealed increased phosphatidylserine (PS) exposure, cell shrinkage and membrane blebbing, delineating an eryptotic phenotype in the host RBC. Interestingly, the blebs on the surface of the iRBCs released to the extracellular milieu become extracellular vesicles (EVs) which are essential for intercellular communication due to their cargo of proteins, nucleic acids, lipids and metabolites. The proteomic characterization displayed 2 highly enriched protein clusters in EVs from CQ-treated iRBCs, the proteasome and ribosome. We demonstrated that this unique protein cargo is not associated with the parasite growth rate. Additionally, we found that these particular EVs might activate IFN signaling pathways mediated by IL-6 in THP-1-derived macrophages. Our findings shed new insights into a novel drug-induced cell death mechanism that targets the parasite and specific components of the infected host RBC.

**IMPORTANCE:** Our previous studies have shown that chloroquine (CQ) treatment in iRBCs triggers *Plasmodium falciparum* digestive vacuole (DV) membrane permeabilization leading to calcium redistribution. Interestingly, increased intracellular calcium concentration is the main inducer of the suicidal death of red blood cells (RBCs) called eryptosis. The present study shed new insights into a novel CQ-induced cell death mechanism that targets the parasite and the infected host RBC by inducing key phenotypic hallmarks of eryptosis: PS exposure, cell shrinkage and membrane blebbing. Moreover, the proteomic characterization of the blebs released to the extracellular milieu also known as extracellular vesicles (EVs) revealed a cargo highly enriched in ribosomal proteins and proteasome subunits relevant for host-parasite interactions. These findings highlight CQ’s effect on calcium homeostasis disruption in infected red blood cells (iRBCs) with cellular and immunological consequences of great significance for malaria pathogenesis and potential clinical implications.

## INTRODUCTION

Malaria is considered one of the most devastating infectious diseases worldwide with an estimated 241 million cases reported in 2020. Of this number, 95% belongs to the WHO African Region having the highest share of the global malaria burden (1). Therefore, efforts towards its control and eradication through novel antimalarial drugs; as well as the search for the long-awaited vaccine is urgently needed. The disease is caused by the *Plasmodium* protozoan, transmitted by the female mosquito of the genus *Anopheles sp.* Currently, it is known that seven species of the parasite are the cause of the disease, *P. falciparum*, *P. vivax*, *P. malariae*, *P. ovale* and the simian species, *P. knowlesi*, *P. cynomolgi* and *P. simium.* Of these, only *P. falciparum* results in high mortality as a result of its prevalence, virulence and drug resistance (1). Malaria symptoms include fever, headache, chills, among others (2), occurring as a consequence of ruptured infected red blood cells (iRBCs) that initiate a systemic release of pro-inflammatory cytokines and other molecules triggering the malaria pathogenesis (3). For this reason, the development of resistance is still an issue for antimalarial compounds targeting the blood stage which leads to its discontinuity for routine treatment in most of the malaria-endemic regions.

Chloroquine (CQ) is a drug that has been used to prevent and treat malaria since the 1940s, however, its ineffective presence has been linked to genetic pressure on erythrocytic-stage parasites resulting in resistance (4). CQ acts on *Plasmodium* late stages (trophozoites and schizonts) by entering the digestive vacuole (DV) to prevent the parasite detoxification process of heme polymerization into hemozoin (5). In 2011, our group demonstrated an alternative mechanism of action happening *in vitro* (6) which was confirmed subsequently by ex *vivo* assays showing its clinical relevance (7). The studies showed that CQ treatment caused permeability of the parasite DV membrane resulting in calcium leakage into the iRBC cytoplasm, involving features of programmed cell death in the parasite, for example, mitochondrial outer membrane permeabilization and DNA degradation. However, no further investigations were pursued on its implications towards the host RBC.

In the past years, it has been reported that an increase in intracellular calcium leads to the suicidal death of RBCs also known as eryptosis (8). Interestingly, during *Plasmodium* invasion and development, the parasite employs a strategy to delay eryptosis allowing its survival by storing calcium in different compartments, one of them being the DV (9). Therefore, we proposed that calcium homeostasis disruption in late-stage iRBCs due to CQ treatment might trigger eryptotic hallmarks. From our data, CQ-treated iRBCs displayed higher levels of phosphatidylserine (PS) externalization and ceramide abundance on the plasma membrane, cell shrinkage, and membrane blebbing; all of them considered classical features of eryptosis.

Following the downstream cellular events of CQ-treated iRBCs, we characterized the membrane blebs released to the extracellular milieu also known as extracellular vesicles (EVs). These are spheres made from a lipid bilayer membrane carrying proteins, metabolites, lipids and nucleic acids from the parasite and the host cell (10). Notably, RBC-derived EVs have been divided into three groups according to their size: Exosome-like vesicles (30 - 100 nm), microvesicles (100 - 1000 nm) and eryptotic bodies (500 - 5000 nm). For their biogenesis, all of them derived directly from the outward budding and fission of the plasma membrane (11). In malaria, EVs are involved in cell-cell communication during the life cycle, immunomodulation, as stage-dependent biomarkers, among others (11). Studies on the roles of EVs in malaria pathogenicity are slowly emerging (12–16), but drug-induced consequences have not yet been established. For this reason, we evaluated their role in parasite invasion and THP-1-derived macrophage stimulation. We found that EVs from CQ-treated iRBCs isolated from *Plasmodium* late stages are highly enriched in proteasome subunits and ribosomal proteins. Based on our findings, we speculate that this distinguished protein profile is not involved in boosting the parasite growth rate. Nevertheless, it might be associated with the activation of IFN signaling pathways mediated by IL-6 in THP-1-derived macrophages with either pro- or anti-inflammatory effects.

Intriguingly, our results shed new insights into a novel CQ-induced cell death mechanism that targets the parasite and specific components of the infected host cell as an exploratory study of the drug’s downstream events during treatment.

## RESULTS

### Chloroquine treatment triggers the externalization of phosphatidylserine and ceramide abundance in the iRBC plasma membrane

To determine if iRBCs expose PS after CQ treatment due to calcium redistribution, synchronized *P. falciparum* 3D7 cultures at 10% parasitemia (mid-late trophozoites) were incubated for 10 hours with 1 μM of Ca^2+^ ionophore (ionomycin), CQ, Mefloquine (MQ) and 1X PBS (non-treated iRBCs). MQ was included as a DV non-disrupting antimalarial drug and uninfected red blood cells (uRBCs) as a negative control due to their low basal levels of PS exposure. Only Ca^2+^ ionophore-treated uRBCs and iRBCs were incubated with malaria culture media (MCM) and ringer solution (RS) to stimulate PS flipping as a positive control. Enriched schizonts by magnetic-activated cell sorting (MACS) were used for Annexin V-FITC and Hoescht staining. The results showed a significant rise of PS exposure for Ca^2+^ ionophore- and CQ-treated iRBCs (Fig. 1a). Furthermore, fluorescence microscopy was used to obtain images from non-treated iRBCs, Ca^2+^ ionophore-treated iRBCs and drug-treated iRBCs after 10 hours of incubation to show PS flipping to the outer leaflet of the plasma membrane which supports the flow cytometry data (Fig. 1b). As reported in the literature, increased intracellular calcium in RBCs triggers the activation of platelet-activating factor (PAF) which stimulates sphingomyelinase to cleave sphingomyelin producing ceramide that accumulates in the plasma membrane. Ceramide abundance was determined by staining MACS-enriched schizonts with anti-ceramide antibody and Hoechst. The results displayed the same trend obtained by PS exposure where Ca^2+^ ionophore-treated and CQ-treated iRBCs had significantly higher levels of ceramide compared to non-treated iRBCs (Fig. 1c). These findings demonstrate that PS exposure and ceramide abundance are exclusive eryptotic features exhibited by CQ-treated iRBCs.

**Figure 1.**
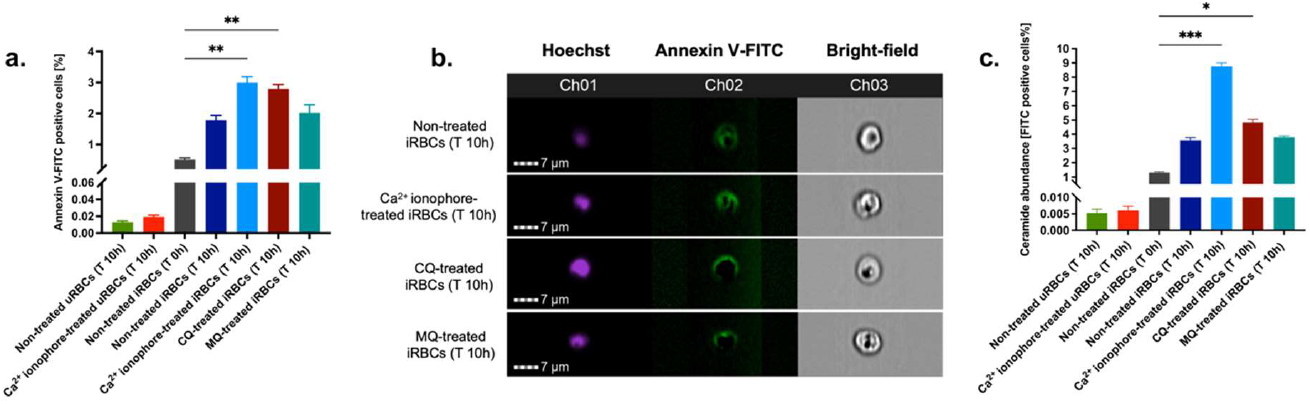
Effect of CQ on PS exposure and ceramide abundance in iRBCs. (a) PS exposure in iRBCs represented by the percentage of Annexin-V FITC positive cells. (b) Fluorescence microscopy images of PS flipping to the outer leaflet of iRBCs plasma membrane. (c) Ceramide abundance in iRBCs represented by the percentage of FITC-positive cells. The results correspond to the mean±SEM of 3 independent experiments; \**p* < 0.05, \*\**p* < 0.01, \*\*\**p* < 0.001 by Kruskal-Wallis test with post-hoc Dunn’s test.

### Digestive vacuole disruption by chloroquine is associated with cell shrinkage and membrane blebbing

Similar to the CQ effect on iRBCs leading to calcium homeostasis disruption and programmed cell death on the parasite, a previous study revealed this novel mode of killing by an obsolete antimalarial drug, Quinacrine (QC) with its ability to induce calcium redistribution at submicromolar levels from DV permeabilization (17). Likewise, a Na^+^/Ca^2+^ exchanger inhibitor, 3’,4’-dichlorobenzamil (DCB) was proposed as a possible new class of DV-destabilizing agent (17). This led us to investigate if CQ, QC and DCB treatment on iRBCs results in an eryptotic morphology such as cell shrinkage and membrane blebbing. Scanning electron microscopy (SEM) was performed using magnet-purified mature parasite forms at the starting point of the treatment and after 10 hours (Fig. 2a). Only QC- and DCB-treated iRBCs showed a significant reduction in the cell diameter, however, the statistical analysis reported a p-value of 0.0556 for CQ-treated iRBCs placing this treatment on the borderline of a significant difference (Fig. 2b). Moreover, the blebs on the surface of iRBCs were counted from 20 scanning electron micrographs per each condition. All drug-treated iRBCs evidenced a significant increase in blebs after 10 hours compared to non-treated iRBCs at T 0h (Fig. 2c). Notably, CQ caused the largest number of blebs from all the compounds tested in the experiment.

**Figure 2.**
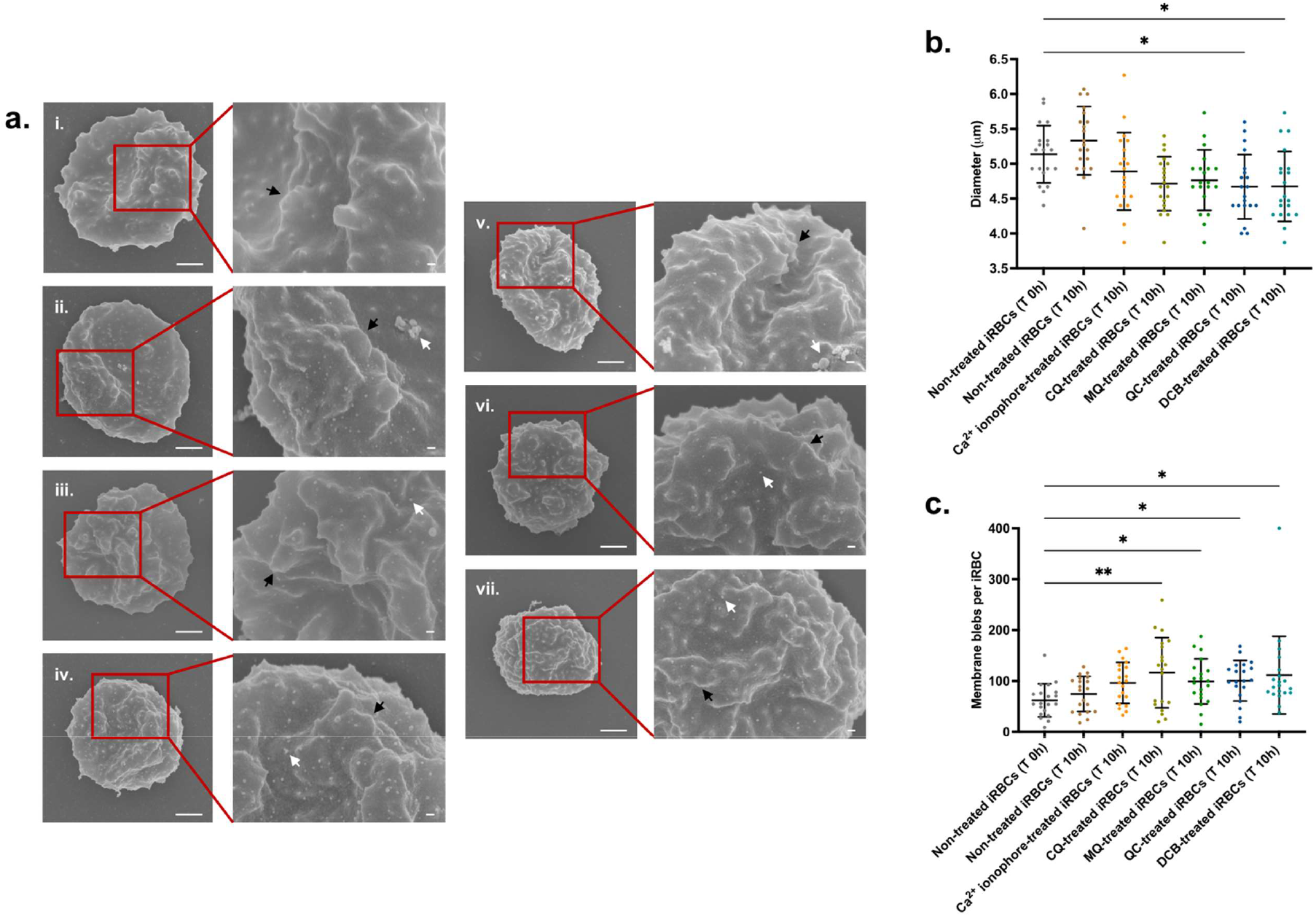
DV-disrupting compounds treatment is associated with an eryptotic morphology in iRBCs: cell shrinkage and microvesiculation. (a) Representative scanning electron micrographs of non-treated iRBCs T 0h (i), non-treated iRBCs T 10h (ii), Ca^2+^ionophore-treated iRBCs T 10h (iii), CQ-treated iRBCs T 10h (iv), MQ-treated iRBCs T 10h (v), QC-treated iRBCs T 10h (vi) and DCB-treated iRBCs T 10h (vii). The black arrows denote knobs and white arrows denote blebs. Scale bars of the original and zoom images represent 1 μm and 100 nm, respectively. (b) Diameter of iRBCs obtained from SEM images. (c) Number of membrane blebs on iRBCs surface quantified from SEM images. For b and c, 20 images per condition were randomly selected and used for the scatter plots. The data represent the mean ± SD; *p < 0.05, **p < 0.01 by Kruskal-Wallis test with post-hoc Dunn’s test.

### Calcium redistribution by chloroquine, quinacrine, and 3’,4’-dichlorobenzamil disrupt the iRBC cytoskeleton network

It has been reported that an increase in intracellular calcium concentration activates calpain, a calcium-dependent cytosolic protease from the host RBC which leads to cytoskeleton cleavage (18). *Plasmodium* exports numerous proteins to the RBC plasma membrane throughout its development, late stages particularly translocate the knob-associated histidine-rich protein (KAHRP) which interacts with the RBC cytoskeleton to form protrusions on the cell surface known as knobs (19). To determine the effects of the antimalarial compounds as RBC cytoskeleton network disruption inducers, μ-calpain activation and KAHRP levels were analyzed by western blot experiments. iRBCs at mid-late trophozoites were treated for 10 hours with CQ, MQ, QC, Artesunate (AS) and DCB. KAHRP bands were quantified using Image J software. The DV-disrupting compounds CQ, QC, and DCB led to doublet bands for μ-calpain (Fig. 3a). In the presence of calcium, the 80 kDa form of μ-calpain is cleaved to the 75 kDa form with increased catalytic activity. Furthermore, these compounds caused a significant loss of KAHRP, whereas the non-disrupting antimalarial drugs MQ and AS did not (Fig. 3a & 3b). To confirm that calcium redistribution is the main factor for this proteolytic processing, a 30 min pre-incubation with the calcium chelator BAPTA-AM followed by a 10-hour treatment with CQ, QC and DCB. The compounds were selected based on their significance displayed previously. The data showed that BAPTA-AM reduced μ-calpain activation in iRBCs treated with CQ, QC and DCB (Fig. 3c). The densitometry plots of μ-calpain were obtained using Image J software to endorse that the second peak corresponding to the 75 kDa form is absent with BAPTA-AM (Figure 3d). Likewise, the calcium chelator pre-treatment demonstrated a significant rescue of KAHRP levels (Fig. 3c & 3e). These findings demonstrate that calcium homeostasis disruption triggered by CQ, QC and DCB treatment in iRBCs mediates calpain activation and loss of KAHRP.

**Figure 3.**
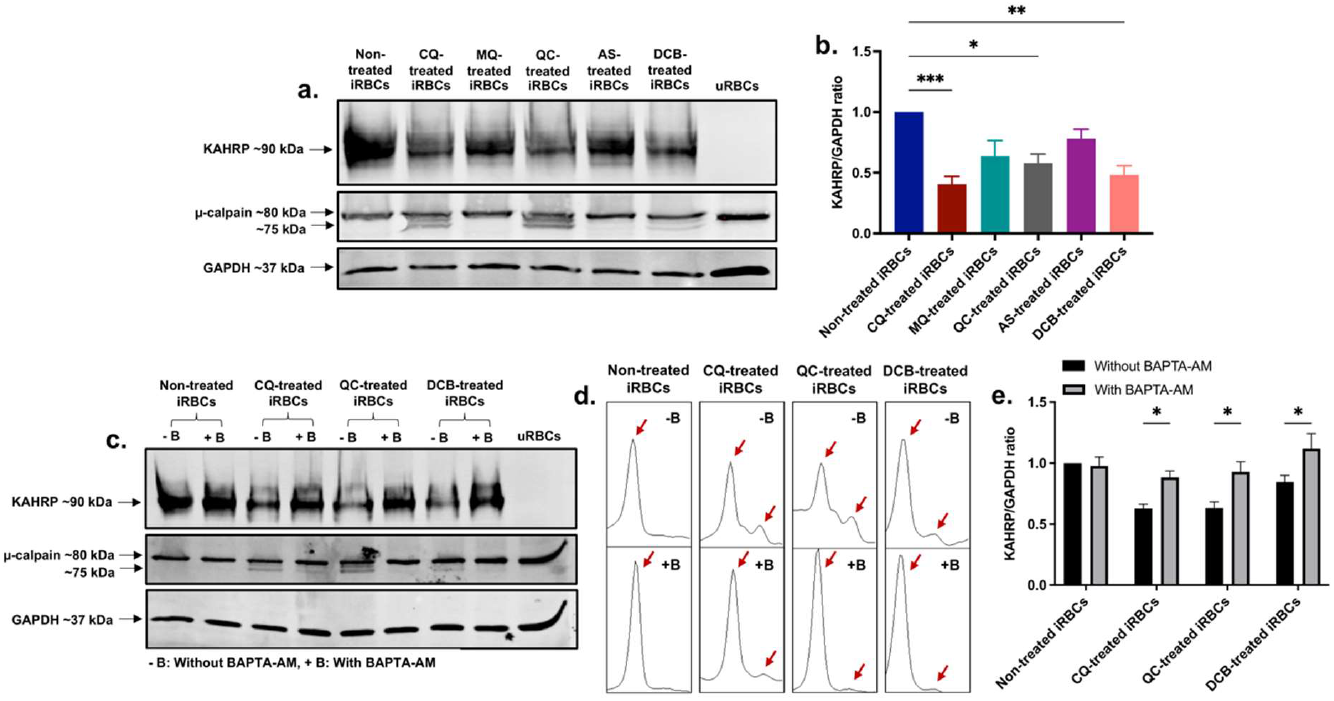
KAHRP degradation and μ-calpain activation are mediated by calcium redistribution in iRBCs treated with CQ, QC and DCB. (a) Western blot of KAHRP and μ-calpain protein expression in drug-treated iRBCs. (b) Quantification of KAHRP/GAPDH ratio normalized to non-treated iRBCs. (c) Western blot of KAHRP and μ-calpain protein expression in drug-treated iRBCs as a result of preincubation with BAPTA-AM. (d) Densitometry plots of μ-calpain bands with and without BAPTA-AM. (e) Quantification of KAHRP/GAPDH ratio normalized to non-treated iRBCs without BAPTA-AM. The data represent the mean±SEM of 6 independent experiments; \**p* < 0.05, \*\**p* < 0.01, \*\*\**p* < 0.001 by Kruskal-Wallis test with post-hoc Dunn’s test (b) and two-way ANOVA test with post-hoc Bonferroni’s test (e).

### iRBCs-derived EVs quantity, distribution and protein profile

The RBC membrane blebs released to the extracellular milieu also named extracellular vesicles (EVs) are important mediators of parasite-parasite and parasite-host communication. EVs from non-treated uRBCs, non-treated iRBCs and CQ-treated iRBCs were isolated by differential ultracentrifugation. Nanoparticle tracking analysis (NTA) was employed to determine the number of EVs (particles/mL) per size ranging from 0 - 1000 nm. Only for 3 size categories EVs from uRBCs were significantly abundant compared to EVs from non-treated and CQ-treated iRBCs (Fig. 4a). As an average size, all EVs samples measured 200 nm approximately (Fig. 4b). Once the EV size was divided into two groups it was observed that non-treated uRBCs preferentially released smaller EVs within the size category 0 - 500 nm, having a mean value of 2.54 × 10^10^ particles/mL (Fig. 4c). However, CQ-treated iRBCs secreted larger EVs within the size category 500.5 - 1000 nm with a mean value of 2.80 × 10^8^ particles/mL (Fig. 4c). EVs were visualized by transmission electron microscopy (TEM), validating the cup-shaped morphology and the presence of lipid bilayer-enclosed nanosized structures (Fig. 4d) which coincides with the average size reported by NTA. Liquid Chromatography Mass Spectrometry (LC-MS) analysis was performed to determine the protein profile of EVs produced under RBC’s physiological condition, parasite development and antimalarial drugs treatment. The set of samples described above in addition to EVs from MQ-treated iRBCs were used. MQ was included in view of its non-disruptive properties on calcium homeostasis from DV membrane permeabilization. Among the proteins identified above the detection threshold, 56 human and 42 parasite proteins were common in all samples (Fig. 4e). Likewise, unique proteins were recognized for each sample. Interestingly, the Kyoto Encyclopedia of Genes and Genomes (KEGG) pathway analysis revealed significant enrichment of human proteasomal proteins in EVs from non-treated iRBCs and EVs from drug-treated iRBCs (Fig. 4f), whereas *Plasmodium* proteasome and ribosome were only significantly enriched in EVs from non-treated iRBCs and EVs from CQ-treated iRBCs (Fig. 4g). These results were relevant to understand that parasite development along with drug pressure might have a role in the size and protein composition of EVs released by iRBCs.

**Figure 4.**
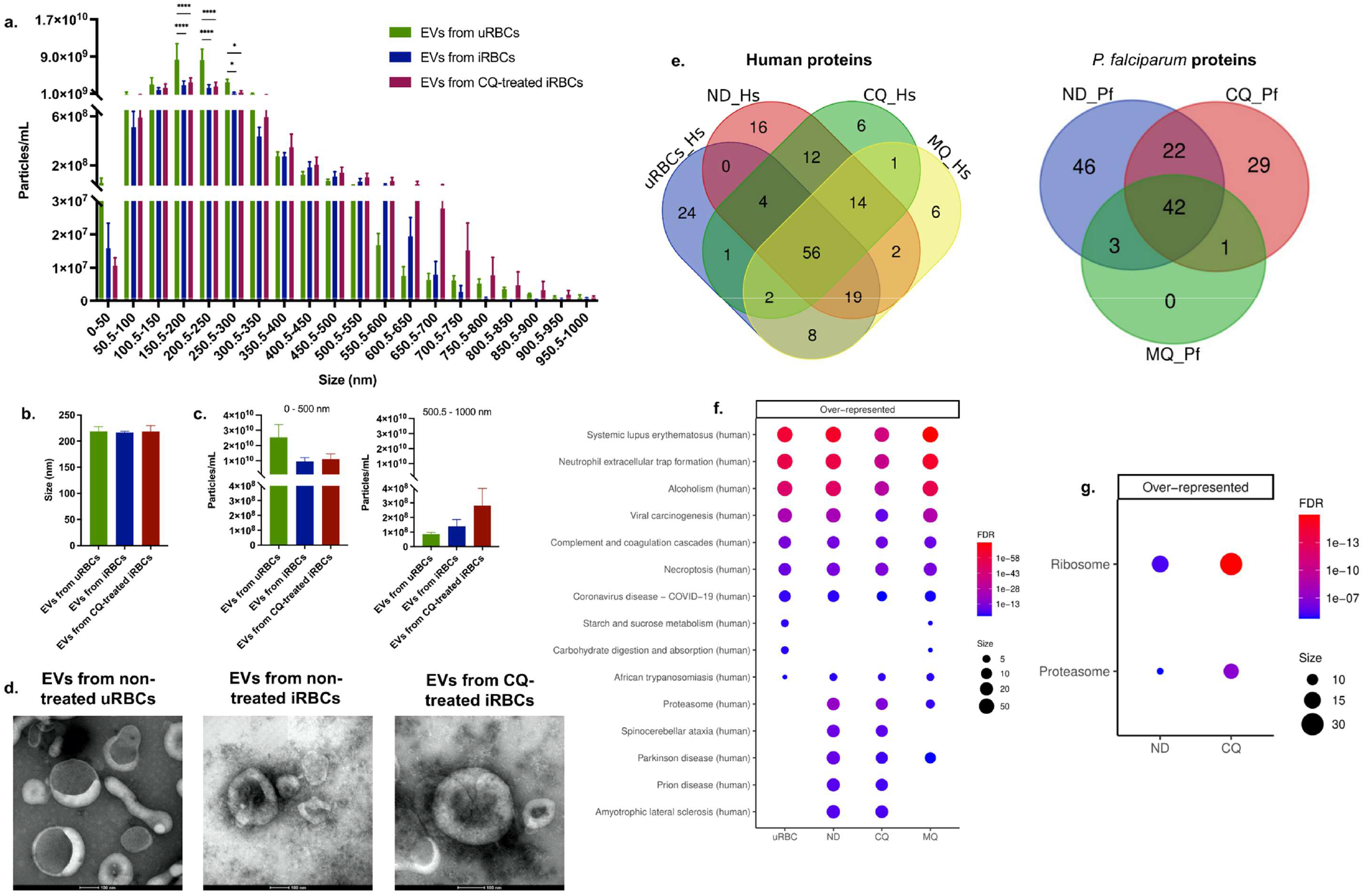
Characterization of iRBCs-derived EVs. (a) Number of EVs from non-treated uRBCs, non-treated iRBCs, and CQ-treated iRBCs per size categories between 0 - 1000 nm obtained by NTA. (b) Particle size average from NTA measurements. (c) Total number of particles, from 0 - 500 nm and 500.5 - 1000 nm. (d) Representative transmission electron micrographs of EV samples (scale bars = 100 nm; original magnification at 35,000×). (e) Venn diagrams illustrating the unique and shared human and parasite proteins identified above the detection threshold in EV samples by LC-MS. KEGG pathway enrichment analysis of (f) human and (g) parasite proteins from EV samples. The results are presented in mean ± SEM of 3 independent experiments; \**p* < 0.05, \*\*\*\**p* < 0.0001 by two-way ANOVA test with post-hoc Bonferroni’s test (a) and Kruskal-Wallis test with post-hoc Dunn’s test (b & c). uRBCs: EVs from non-treated uRBCs; ND: EVs from non-treated iRBCs; CQ: EVs from CQ-treated iRBCs; MQ: EVs from MQ-treated iRBCs.

### Chloroquine treatment leads to an enrichment of proteasome subunits and ribosomal proteins in iRBCs-derived EVs

In order to quantify the differential abundance of EV proteins in response to antimalarial drug treatment, we used Sequential Window Acquisition of all Theoretical Mass Spectra (SWATH-MS) analysis with several statistical cut-off criteria, for example, a minimum of two unique peptides quantified per protein, fold change ≥ 1.5 and adjusted p-value ≤ 0.05. Protein-protein interaction networks by STRING analysis highlighted the 4 main clusters of the entire human and *Plasmodium* proteome found in our samples (Fig. 5a and 5d). Intriguingly, the proteasome protein cluster was found for human and Plasmodium datasets. The relative abundance of the 50 most abundant proteins of each dataset was visualized in heatmaps. For example, hemoglobin subunits alpha and beta were highly enriched as expected in EVs from non-treated uRBCs due to the absence of parasite invasion (Fig. 5b). Meanwhile, ribosomal proteins and proteasome subunits were significantly enriched in EVs from CQ-treated iRBCs compared to EVs from non-treated and EVs from MQ-treated iRBCs (Fig. 5e). Gene ontology (GO) analysis was performed to obtain functional insights on the EV protein cargo. For human proteins, no pathways associated with the proteasome complex were found in cellular component, biological process and molecular function (Supplemental Fig. 1a-c). On the other hand, parasite proteins displayed proteasome core complex and proteasome alpha-subunit complex as strongly enriched cellular components in EVs from CQ-treated iRBCs (Supplemental Fig. 1d and Supplemental Table 1). KEGG analysis determined the enriched pathways in both datasets. The proteasome pathway was upregulated for human (Fig. 5c) and parasite (Fig. 5f) proteins in EVs from CQ-treated iRBCs, whereas the parasite ribosome pathway (Fig. 5f) was upregulated in EVs from drug-treated iRBCs (Supplemental Table 1). Furthermore, GSEA plots of the top 3 significant KEGG pathways from parasite proteins in EVs from CQ-treated iRBCs and EVs from MQ-treated iRBCs compared with each other and to EVs from non-treated iRBCs are shown in Fig. 5g. Similarly, cell lysates were assessed by SWATH-MS where GO and KEGG analysis indicated human proteasome-related pathways and parasite ribosome-related pathways to be upregulated in CQ-treated iRBCs in contrast with non-treated iRBCs (Supplemental Fig. 2). These results delineate that proteasome and ribosome protein clusters are characteristic phenotypes of EVs from CQ-treated iRBCs.

**Figure 5.**
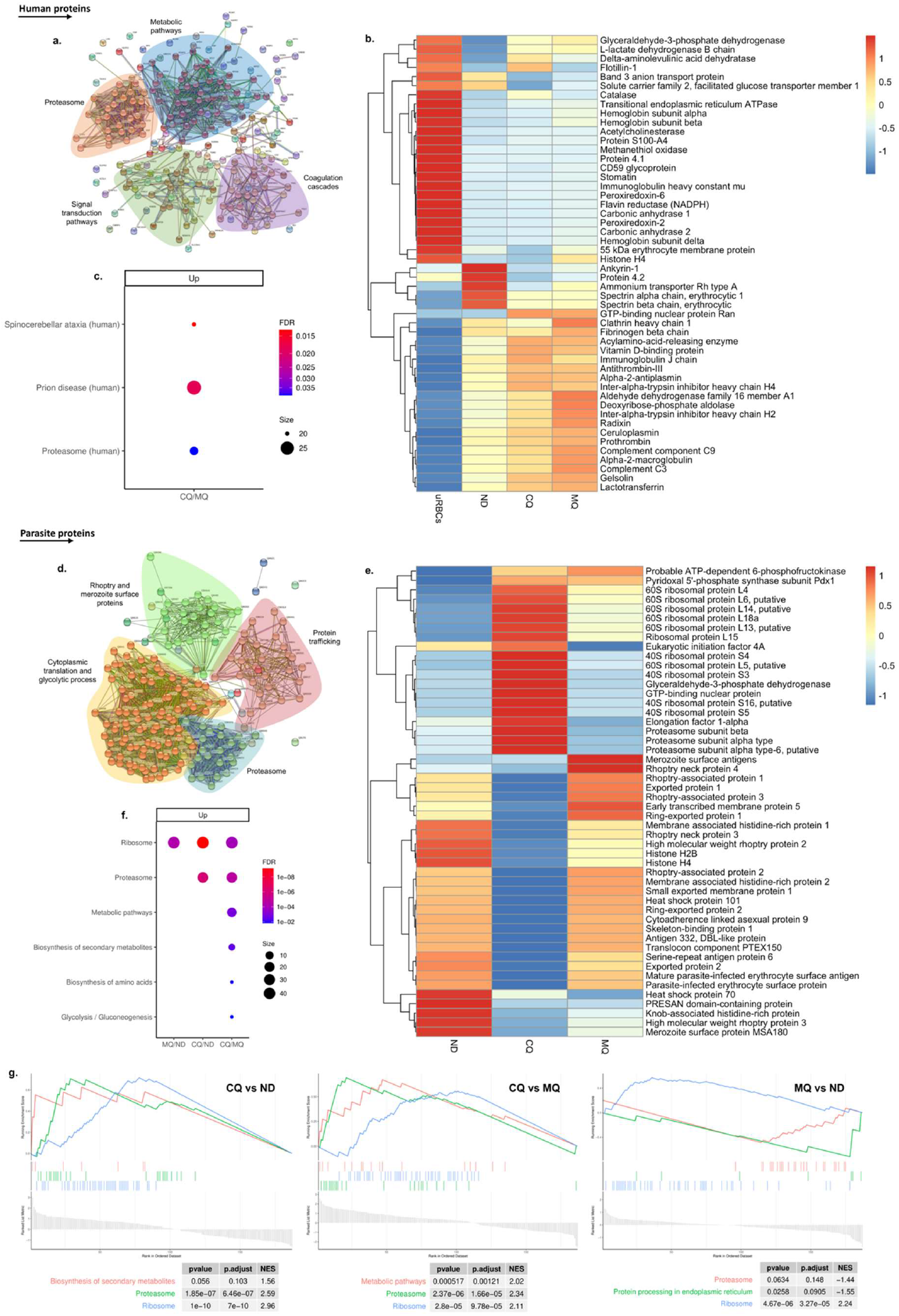
Quantitative proteomic data of EVs. (a) Human and (d) parasite protein interaction networks of EV samples. Whole proteins identified from both datasets were used. Protein nodes and interactions were coloured according to MCL clustering using STRING software. Heatmap of the 50 most abundant (b) human and (e) *P. falciparum* proteins. The average of 3 independent experiments per sample is shown. The colours represent Z score mean protein expression values for samples. KEGG enrichment analysis of (c) human and (f) parasite proteins. Bubble size: Number of proteins assigned to the pathway; Colour: adjusted p-value. (g) GSEA analysis of the top 3 positively and negatively enriched KEGG pathways from parasite proteins. uRBCs: EVs from non-treated uRBCs; ND: EVs from non-treated iRBCs; CQ: EVs from CQ-treated iRBCs; MQ: EVs from MQ-treated iRBCs.

Differentially expressed proteins from both datasets were indicated in volcano plots. 6 human proteasomal subunits were upregulated in EVs from CQ-treated iRBCs compared to EVs from non-treated and EVs from MQ-treated iRBCs (Fig. 6a & Supplemental Table 2). 23 and 16 parasite ribosomal proteins were upregulated in EVs from CQ-treated iRBCs compared to EVs from non-treated and EVs from MQ-treated iRBCs, respectively (Fig. 6b & Supplemental Table 3); 13 were shared between both conditions (Fig. 6c). 8 and 13 parasite proteasome subunits were upregulated in EVs from CQ-treated iRBCs compared to EVs from non-treated and MQ-treated iRBCs, respectively (Fig. 6b & Supplemental Table 3); 8 were shared between both conditions (Fig. 6d). Furthermore, some of them were identified exclusively in one condition. Although CQ and MQ are quinoline-based compounds that inhibit hemozoin formation in the parasite DV, only CQ is capable to trigger DV permeabilization-induced calcium dysregulation which might be associated with its unique EV cargo.

**Figure 6.**
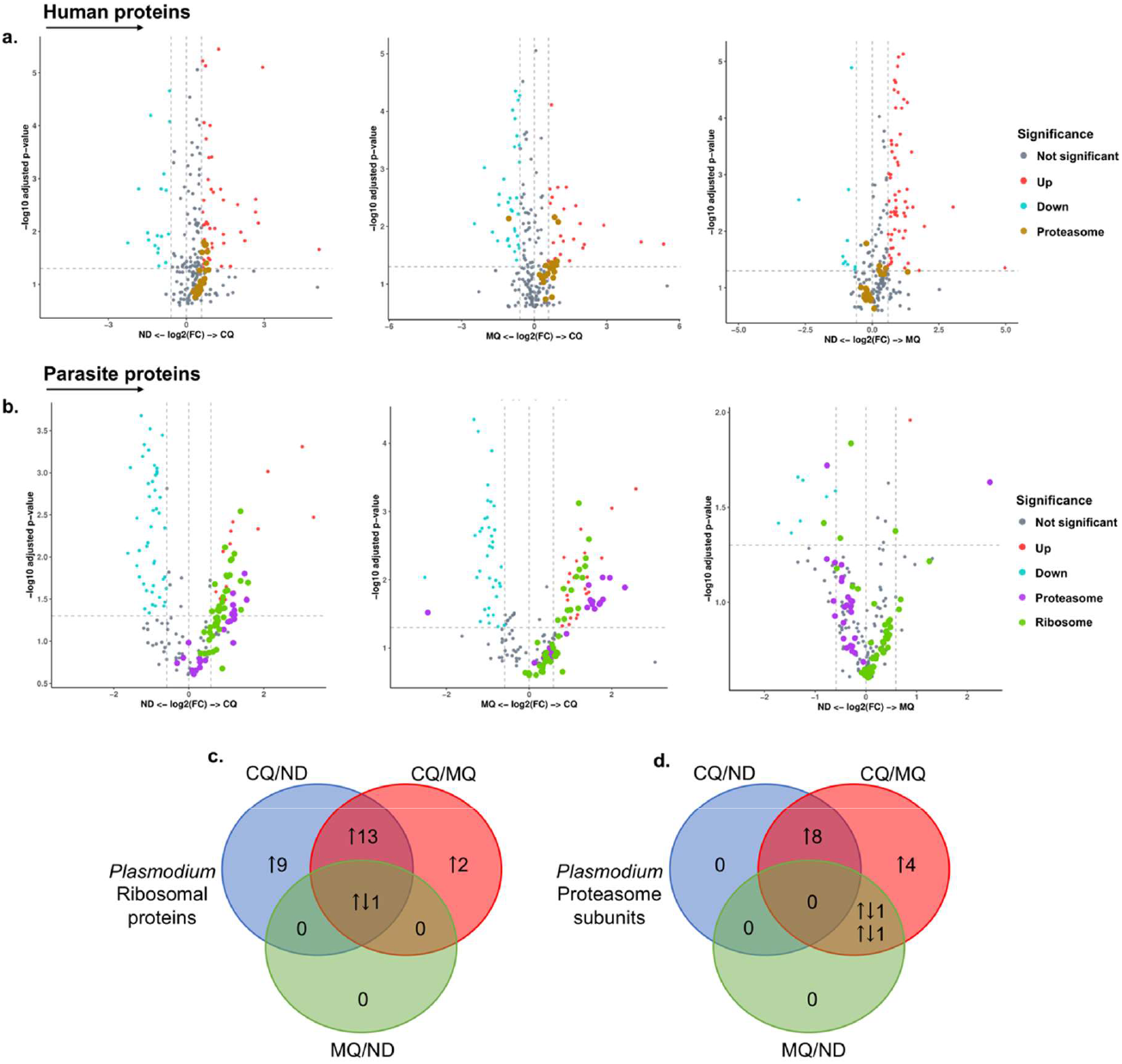
Significantly enriched proteasome and ribosomal proteins in iRBCs-derived EVs. (a) Volcano plots of differentially expressed human proteins in iRBCs-derived EVs. Proteasome subunits (brown dots) are highlighted. (b) Volcano plots of differentially expressed *P. falciparum* proteins in iRBCs-derived EVs. Proteasome subunits (purple dots) and ribosomal proteins (green dots) are highlighted. X-axis, vertical-dotted lines: log2(FC) cut-off is 0.585, corresponding to fold change = 1.5. Y-axis, horizontal-dotted line: Significance cut-off, adjusted p-value = 0.05 (a & b). (c) Venn diagram of significantly differentially expressed parasite ribosomal proteins. (d) Venn diagram of significantly differentially expressed parasite proteasome subunits. The arrows indicate upregulated (↑) and downregulated (↓) proteins (c & d). ND: EVs from non-treated iRBCs; CQ: EVs from CQ-treated iRBCs; MQ: EVs from MQ-treated iRBCs.

### The role of EVs from CQ-treated iRBCs in parasite invasion and THP-1-derived macrophage response

Our results for LC-MS and SWATH-MS have demonstrated a unique protein profile composed of highly enriched ribosomal proteins and proteasome subunits in EVs from CQ-treated iRBCs which might play an essential role in recipient cells. First, we investigated their effect on the parasite growth rate of *P. falciparum* cultures. A 32-hour invasion assay was performed with Cell Tracker Deep Red (CTDR)-labeled uRBCs, MACS-enriched schizonts, and EVs from non-treated uRBCs, EVs from non-treated iRBCs, EVs from CQ-treated iRBCs and 1X PBS (control). Giemsa smears were prepared to determine the parasitemia by light microscopy (Supplemental Fig. 3a), whereas the DNA-specific dye Hoechst was used to determine parasite invasion by flow cytometry. No significant difference in the parasite growth rate was found among EV samples compared to the control for both methods (Supplemental Fig. 3b). This result suggests that the high content of proteasomal subunits and ribosomal proteins in EVs from CQ-treated iRBCs do not play a role in the invasion efficiency.

The effect of EVs on the host’s innate immune response was studied *in vitro* using THP-1-derived macrophages. THP-1 cells were differentiated with phorbol 12-myristate 13-acetate (PMA) into macrophages and stimulated with EVs from non-treated uRBCs, EVs from non-treated iRBCs and EVs from CQ-treated iRBCs for 14 hours. LPS and media were used as a positive and negative control, respectively. The cell culture supernatant was collected for ELISA to measure TNF, IL-6 and IL-1β levels, and cells were collected for transcriptomic analysis. EVs from CQ-treated iRBCs significantly induced IL-6 production although the effect was not as strong as LPS (Supplemental Fig. 4). Meanwhile, TNF and IL-1β have not significantly changed. Transcriptomic profiling showed that EVs from non-treated uRBCs, EVs from non-treated iRBCs and EVs from CQ-treated iRBCs induced 23, 125 and 78 differentially expressed genes (DEG), respectively (Fig. 7a). The top 40 DEGs for EVs from CQ-treated iRBCs against media were displayed in a heatmap (Fig. 7b), revealing some genes with a similar relative expression in cells stimulated with EV samples. Interestingly, the Reactome pathway analysis indicated interleukin pathways as over-represented among DEGs for THP-1-derived macrophages stimulated with EV samples (Fig. 7c). Whereas interferon signaling and interferon alpha/beta signaling were unique over-represented pathways upon stimulation with EVs from CQ-treated iRBCs. For KEGG analysis, TNF signaling pathway, cytokine-cytokine receptor interaction and viral protein interaction with cytokine and cytokine receptor were over-represented pathways for THP-1-derived macrophages stimulated with EV samples (Fig. 7d). Additionally, GO analysis disclosed biological process pathways associated to viral life cycle for THP-1-derived macrophages stimulated with EVs from non-treated and EVs from CQ-treated iRBCs (Supplemental Fig. 5b). DEGs of two pathways were featured in volcano plots (Fig. 7e). The interferon alpha/beta signaling from Reactome pathways exhibited 3 interferon stimulated genes (ISG) upregulated only for EVs from CQ-treated iRBCs, while cytokine-cytokine receptor interaction from KEGG pathways displayed 4, 11, 8 DEGs for EVs from non-treated uRBCs, EVs from non-treated iRBCs and EVs from CQ-treated iRBCs, respectively (Supplemental Table 4).

**Figure 7.**
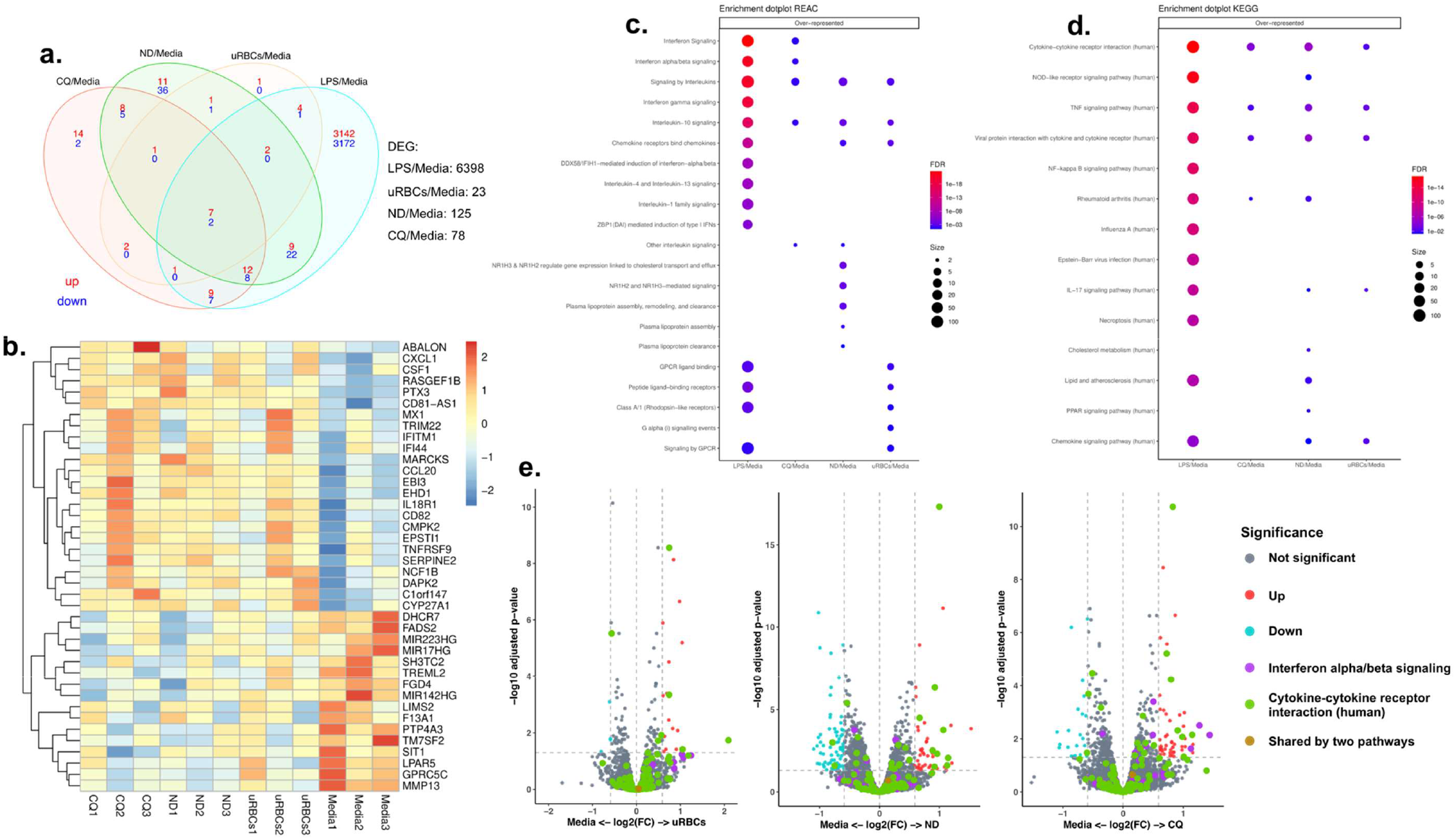
RNA sequencing data of THP-1-derived macrophages upon stimulation with EVs. (a) Venn diagram of significantly DEG. Upregulated and downregulated genes are shown in red and blue, respectively. (b) Heatmap of top 40 DEG from THP-1-derived macrophages stimulated with EVs from CQ-treated iRBCs. The colours represent Z score mean protein expression values for samples. Over-representation of (c) Reactome and (d) KEGG pathway analysis. Bubble size: Number of genes assigned to a pathway; Colour: adjusted p-value. (e) Volcano plots denote the upregulated and downregulated genes of Reactome (purple dots) and KEGG (green dots) pathways. X-axis, vertical-dotted lines: log2(FC) cut-off is 0.585, corresponding to fold change = 1.5. Y-axis, horizontal-dotted line: Significance cut-off, adjusted p-value = 0.05. uRBCs: EVs from non-treated uRBCs; ND: EVs from non-treated iRBCs; CQ: EVs from CQ-treated iRBCs.

## DISCUSSION

Our previous findings have demonstrated that CQ leads to selective permeabilization of the *P. falciparum* DV and mitochondria membrane triggering calcium leakage to the iRBC cytoplasm. Here, we have presented evidence that CQ-induced calcium homeostasis disruption is associated with eryptotic hallmarks in the host RBC and a distinct EV proteomic profile.

PS flipping is considered one of the main features during eryptosis (9). We assessed the impact of CQ treatment in mid-late trophozoites iRBCs and observed a higher exposure of PS. CQ’s effect on intracellular calcium dysregulation may induce the calcium-dependent scramblases that move PS facing outward in the plasma membrane. Furthermore, the literature provides evidence that increasing levels of intraerythrocytic calcium have been associated with higher levels of platelet-activating factor that stimulates sphingomyelinase to cleave sphingomyelin (20). Consequently, ceramide is produced and accumulated in the plasma membrane causing lipid scrambling and PS exposure (21, 22). Our data agree with the previous findings demonstrating that ceramide levels significantly increase in CQ-treated iRBCs.

In 2014, our group demonstrated that calcium redistribution induced by CQ, QC and DCB leads to programmed cell death features in parasite cultures such as mitochondrial depolarization and DNA loss (17). However, no further research was done on the downstream effects of these deleterious compounds in the host RBC. Our current findings have shown that CQ, QC and DCB caused cell shrinkage in iRBCs. As reported in the literature, an increase in intracellular calcium activates the calcium-sensitive potassium channels (Gardos) which prompt iRBC membrane hyperpolarization by potassium and chlorine efflux with osmotic dehydration resulting in cell shrinkage (9, 23). Furthermore, SEM images showed a significant number of blebs on the surface of drug-treated iRBCs being the most abundant CQ-treated iRBCs. The data was confirmed by the activation of the calcium-dependent protease μ-calpain which cleaves the RBC cytoskeleton leading to protein network disruption in the plasma membrane after CQ, QC and DCB treatment. Interestingly, μ-calpain also known as calpain-1 has been reported to negatively regulate RBC deformability and filterability without any visible effect on RBC lifespan under physiological conditions (24). By using a calpain-1 knockout mouse model, calpain-1 function was directly involved in the RBC shape regulation and the shape transition rate in a calcium-dependent manner (24). Along with our data, we propose that μ-calpain participates actively in cytoskeletal and membrane integrity disruption in iRBCs. Furthermore, western blot experiments evidenced that the knob protein KAHRP was less abundant and its rescue together with a decrease of the μ-calpain catalytic subunit was associated with the presence of the calcium chelator BAPTA-AM. Therefore, membrane blebbing is part of the catastrophic events associated with calcium homeostasis disruption by CQ, QC and DCB treatments in iRBCs.

Following the downstream effects of CQ treatment on iRBCs, we isolated and characterized the blebs released to the extracellular milieu as EVs to explore their phenotype. It has been demonstrated that malaria EVs have an average diameter of 200 nm with an intact lipid bilayer (12, 14, 25, 26). Likewise, our TEM and NTA data agreed with these studies. We classified the number of EVs per size category to determine small and larger groups. EVs from uRBCs were predominantly smaller and quite abundant. Previous studies stated that RBCs stored above 2 weeks released significantly more EVs with high protein levels which proportionally increase with the age of the cell (27–29). For this study, we worked with RBCs within the first 3 weeks after collection. In addition, delivery time and cold chain fluctuations might be add-on factors affecting RBCs with a surge in microvesiculation but still supporting the healthy growth of malaria parasites (28, 29). In malaria, during schizont maturation and egress of merozoites, high EV levels were reported which coincide with loss of membrane rigidity and proteolytic disassembly of the RBC cytoskeleton (13). Interestingly, the concentration of EVs represented per size category showed a clear tendency of larger EVs from CQ-treated iRBCs denoting that the cell could be undergoing eryptosis and releasing eryptotic bodies (30).

Proteomic profiling and significant markers from EVs were determined by LC-MS. We found a strong correlation between our protein dataset and the iRBCs-derived EV proteins reported by previous studies (14, 15). Asides from that, we identified the presence of some human and parasite proteins which were exclusive from our EV samples. Therefore, we consider that the characterization outcome of malaria EVs might differ in terms of morphology; size; concentration; nucleic acids, lipids, proteins and metabolites cargo due to the following factors: Parasite stage, culture conditions, EVs isolation method, the solution used to resuspend isolated EVs as well as storage time and temperature.

KEGG pathway analysis of LC-MS datasets revealed that parasite ribosomal proteins along with human and parasite proteasome subunits were found in EVs from non-treated and CQ-treated iRBCs. These results broached our next question to determine if CQ treatment plays a role in the abundance of iRBCs-derived EV proteins. Using SWATH-MS quantitative proteomics we determined that proteasome was one of the main clusters for human and parasite proteins within the EVs. The proteasome is a major degradation machinery essential to preserve proteostasis by irreversibly removing misfolded, damaged or short-lived regulatory proteins (31). The 26S proteasome is ATP-dependent and requires ubiquitin-tagged substrates, meanwhile, the 20S proteasome is ubiquitin- and ATP-independent (32). Interestingly, proteasome subunits have been identified in EVs secreted by other protozoa, helminths and ectoparasites (33). For example, in the protozoa *Acanthamoeba castellanii* the proteasome is involved in the regulation of encystation and extracellular proteolytic activities (34); whereas in *Trichinella spiralis* the proteasome subunit beta type-7 (PST) has been expressed at different developmental stages and is responsible for a decrease of 45.7% adult worm burden in mice suggesting PST as vaccine antigen against the parasite infection (35). Recently, it has been demonstrated that iRBCs-derived EVs contain functional 20S proteasome complexes (15). Our quantitative proteomic data aligns with this finding and provides evidence for the first time, about the upregulation of human and parasite 20S proteasome subunits and human 26S proteasome subunits in EVs from CQ-treated iRBCs. *Plasmodium* 40S and 60S ribosomal proteins were also upregulated in EVs from CQ-treated iRBCs which might be related to ribosome biogenesis disturbance by CQ treatment. In agreement with the literature, chemical agents, lack of nutrients and gene deregulation trigger ribosomal stress which leads to the accumulation of ribosome-free ribosomal proteins (36). Our group has described the detrimental consequences of CQ in iRBCs, precisely in the calcium homeostasis disruption that might be an internal stimulus for downstream mechanisms causing the degradation of the proteasome and ribosome complexes. This represents an important gap in knowledge that requires further investigation.

We investigated the role of EVs from CQ-treated iRBCs in the parasite growth development of *P. falciparum* due to their protein profile enriched in proteasome subunits and ribosomal proteins. Previously, Dekel et al. identified active 20S proteasome complexes along with kinases delivered by iRBCs-derived EVs to uRBCs leading to cytoskeleton network remodeling and altering membrane stiffness for a successful parasite invasion (15). Unlike this finding, we did not observe a significant change in the parasite growth rate in response to EV samples. On the other hand, Vimonpatranon et al. isolated EVs from four *P. falciparum* parasite strains including 3D7 and performed invasion assays to determine the parasite growth development in two invasion cycles (16). 3D7 was the only strain without a significant effect on the invasion efficiency, besides no parasite proteasome was found in the EV cargo. These discrepancies could be explained due to the parasite stage-dependent EV cargo that in both studies were from trophozoites. On the contrary, we used a 10-hour incubation to collect the EVs, starting from mid-late trophozoites (36 - 38 hours p.i.) to schizonts (46 - 48 hours p.i). Also, we calculated the EVs for invasion assays according to the protein yield, in this case, 300 μg representing approximately 4 × 10^9^ particles/mL. Meanwhile, Dekel et al. and Vimonpatranon et al. used EVs based merely on the number of particles, 1 × 10^12^ and 1 × 10^9^ particles/mL, respectively (15, 16). These results remark that the role of EVs in parasite invasion is highly dependent on their cargo and the number of particles used.

EVs in malaria have been described to participate in immunomodulation (11, 37). As it was mentioned before, parasite molecules within EVs vary depending on parasite developmental stages which might impact the immune system differently. Monocytes, macrophages and NK cells response to iRBCs-derived EVs from ring and trophozoite stage have been evaluated (14–16, 38), however, there are no reports using EVs released from the tight timeframe of mid-late trophozoites to schizonts and in the presence of drug pressure. For this reason, we used THP-1-derived macrophages stimulated with these particular EVs. From the proinflammatory cytokines evaluated in our study by ELISA, only a significant release of IL-6 was found as a response to EVs from CQ-treated iRBCs. It has been demonstrated that IL-6 induces the expression of ISGF3γ, a subunit of the transcription factor ISGF3, which in turn promotes IFN responses (39). Intriguingly, we found that the Reactome pathway of interferon alpha/beta signaling was over-represented after stimulation with EVs from CQ-treated iRBCs with ISGs related to antiviral response. These results might hint that THP-1-derived macrophages stimulated with EVs from CQ-treated iRBCs secrete IL-6 which modulates a type I IFN signaling by the expression of ISGs.

Considering the unique protein cargo of EVs from CQ-treated iRBCs previously described, we found that one of the most significantly abundant proteins was *P. falciparum* 60S acidic ribosomal protein P2 (P2). Notably, a recent study using a murine model revealed the immunogenic properties of P2 with high IgG titers after the second immunization which was comparable to the levels obtained by the parasite-specific antigen, Msp-119 (40). Furthermore, experimental evidence has demonstrated that oligomeric/tetrameric P2 is localized on the iRBC surface during late trophozoite/early schizont stage and blocking its accessibility by monoclonal antibodies led to nuclear division arrest of the parasite (41). In this context, P2 together with more ribosomal proteins may have extra-ribosomal roles in iRBCs and contribute as immunogenic antigens during malaria pathogenesis. Based on our previous speculation, functional experiments involving the interaction of parasite ribosomal proteins with immune cells are required to determine if they contribute to a type I IFN response.

From our findings, we have identified some limitations for malaria EVs interaction with RBCs and THP-1-derived macrophages: (a) The activity of the 20S proteasome complex was not evaluated for the RBC cytoskeleton proteins. (b) Lack of transcriptomic and lipidomic analysis of EVs to determine if these molecules had different profiles as a consequence of the drug pressure which might play a role in recipient cells. Therefore, we cannot attribute our results entirely to the parasite proteins. (c) Inhibitory assays for ribosomal proteins and proteasome subunits are required to assure their role in parasite invasion and activation of THP-1-derived macrophages. (d) Timepoint multiplex cytokine assays are needed to determine if EVs from CQ-treated iRBCs trigger a proinflammatory or anti-inflammatory type I IFN response and its progression over time.

Here we are reporting for the first time that CQ-induced calcium redistribution in iRBCs triggers eryptotic hallmarks in the host RBC for a rapid killing effect with downstream events that might be considered a double-edged weapon for parasite pathogenicity (Fig. 8).

**Figure 8.**
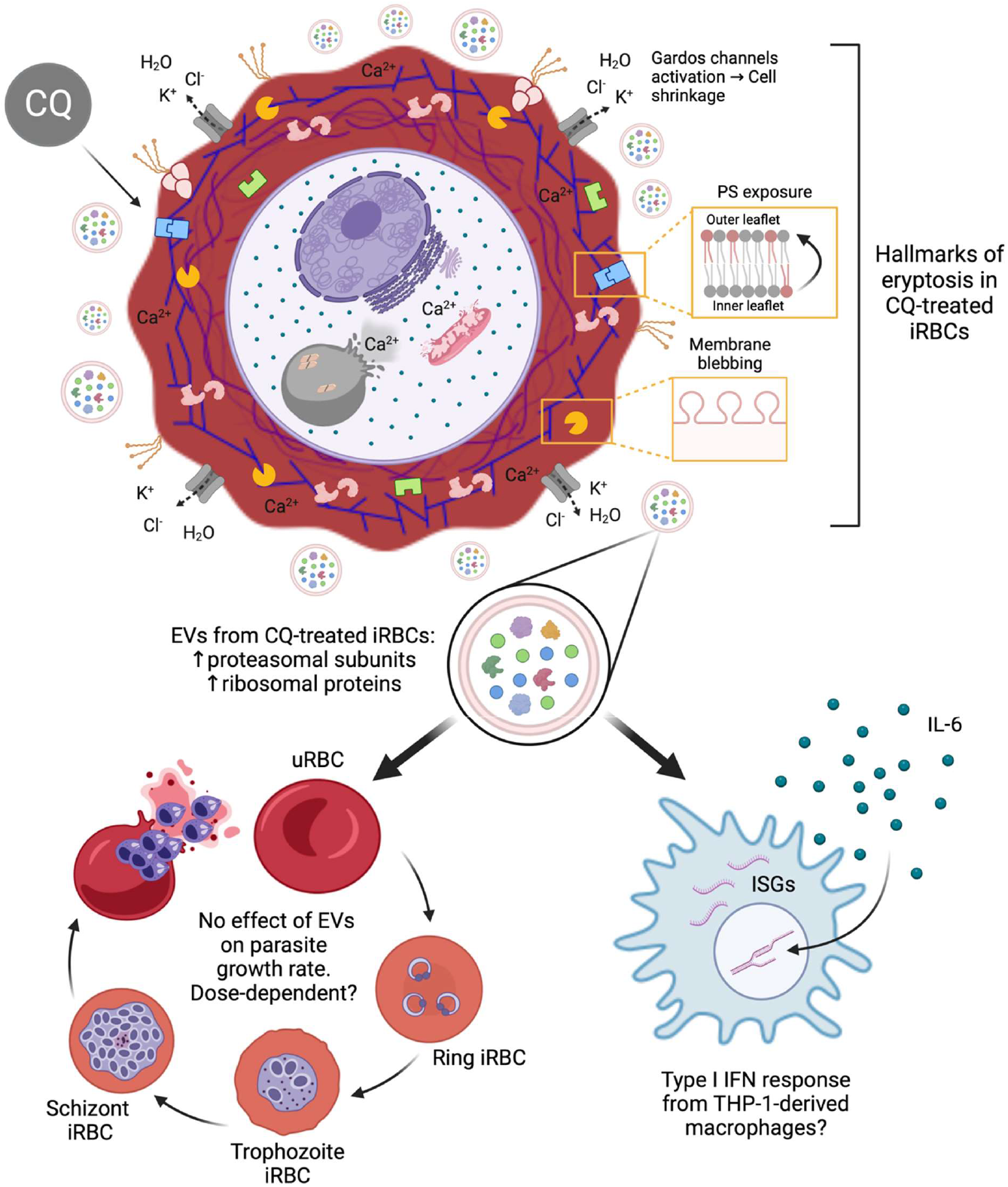
Proposed model of downstream effects triggered by CQ treatment in iRBCs. The diagram was created based on the results observed in this study associated with intracellular calcium homeostasis dysregulation in CQ-treated iRBCs. *Created with BioRender.com*

## MATERIALS AND METHODS

### *Plasmodium falciparum* culture

3D7 *P. falciparum* laboratory strain (MRA-104; MR4, ATCC Manassas Virginia) was cultured and maintained in 25 cm^2^ or 75 cm^2^ non-vented cell culture flasks at 10% parasitemia in 2.5% haematocrit with malaria culture medium (MCM), consisting of a filter-sterilized homogeneous mixture of RPMI 1640 (Life Technologies, USA), 2.2 g/L sodium bicarbonate, 0.5% AlbuMAX II (Gibco, Thermo Fisher Scientific, USA), 0.005% hypoxanthine (Sigma-Aldrich, MO, USA), 0.03% L-glutamine (Sigma-Aldrich, MO, USA), and 25 μg/L gentamicin (Gibco, Thermo Fisher Scientific, USA). The pH was adjusted to 7.4. Type O-positive human RBCs in citrate-phosphate-dextrose with adenine (CPDA-1) anticoagulant were obtained from the Interstate Blood Bank (Memphis, TN, USA) and tested negative for infectious agents according to FDA guidelines. Parasite cultures were incubated under a gas mixture of 5% CO_2_, 3% O_2_, and 92% N2 at 37 °C in the dark. Parasitemia and stage of the culture were assessed by light microscopy after staining methanol-fixed thin films with a diluted Giemsa stain. Parasites were synchronized to select ring-stage parasites with 5% sorbitol (Merck, Germany) at 37°C for 15 min. P. falciparum cultures were tested for mycoplasma once a month using MycoStrip^™^ (InvivoGen, USA).

### Solutions and drug preparation

The positive controls were incubated with the calcium ionophore ionomycin at 2.5% haematocrit in MCM and ringer solution (RS) in a 1:1 ratio. RS was prepared with 125 mM NaCl, 5 mM KCl, 1 mM MgSO_4_, 32 mM N-2-hydroxyethylpiperazine-N-2-ethanesulfonic acid (HEPES), 5 mM glucose, and 1 mM CaCl_2_; the pH was adjusted to 7.4 and the temperature kept at 37°C. All drug compounds (CQ diphosphate, MQ hydrochloride, QC dihydrochloride, AS and DCB hydrochloride) and ionomycin were purchased from Sigma-Aldrich. CQ was diluted in sterile 1× phosphate-buffered saline (PBS), while the remaining drugs and ionomycin were dissolved in sterile dimethyl sulfoxide (DMSO) to stock aliquots of 10 mM, frozen at −20°C and wrapped in aluminium foil. Before any experiment, fresh working drug concentrations were prepared after dilution with sterile 1× PBS to ensure a final assay mixture of less than 0.5% DMSO.

### Annexin V-binding to determine phosphatidylserine exposure

Parasites were synchronised for 2 cycles prior to the start of the experiment. Using 25 cm^2^ flasks, malaria cultures were treated for 10 hours with a final concentration of 1 μM of drug or 1x PBS (control) at 10% parasitemia (mid-late trophozoites) in 2.5% haematocrit. uRBCs cultures were treated only with ionomycin and 1x PBS. Afterward, late stages (schizonts) were enriched using the QuadraMACS Separator and LD columns (Miltenyi Biotec, Germany). iRCBs were stained with Annexin V-FITC (BioVision Inc, USA) in RS at room temperature (RT) for 30 min in the dark. The pellet was stained with Hoescht 33342 (Sigma-Aldrich, USA) for 10 min at a final concentration of 1 μg/ml in the dark. 1x PBS was used to wash the cells twice after each staining step. Annexin V-FITC and Hoescht 33342 fluorescence intensity were measured on a BD LSR Fortessa X-20 (BD, USA).

### Ceramide abundance at RBC surface

Parasites were synchronised for 2 cycles prior to the start of the experiment. Using 25 cm^2^ flasks, malaria cultures were treated for 10 hours with a final concentration of 1 μM of drug or 1x PBS (control) at 10% parasitemia (mid-late trophozoites) in 2.5% haematocrit. uRBCs cultures were treated only with ionomycin and 1< PBS. Afterward, late stages (schizonts) were enriched using the QuadraMACS Separator and LD columns (Miltenyi Biotec, Germany). iRBCs were stained for 1 hour at 37°C with 1 μg/mL ceramide monoclonal antibody (clone MID 15B4) (Enzo Life Sciences, Germany) diluted 1:10 in RS containing 0.1% bovine serum albumin (BSA) and then washed twice using RS-BSA 0.1%. After that, cells were stained for 30 min with polyclonal fluorescein-isothiocyanate (FITC)-conjugated goat anti-mouse IgG H&L specific antibody (Abcam, United Kingdom) diluted 1:50 in RS-BSA 0.1%. The unbound secondary antibody was removed by repeated washing with RS-BSA 0.1%. The pellet was stained with Hoescht 33342 (Sigma-Aldrich, USA) for 10 min at a final concentration of 1 μg/ml in the dark followed by repeated washing with 1 × PBS. FITC and Hoescht 33342 fluorescence intensity were measured on a BD LSR Fortessa X-20 (BD, USA).

### Scanning electron microscopy (SEM)

Images were analyzed with a Nikon Eclipse E800 microscope (Nikon). For SEM, iRBCs coated on poly-lysine (Sigma-Aldrich) glass coverslips were fixed in 2.5% glutaraldehyde, washed, treated with 1% osmium tetroxide (Ted Pella), and critical point dried (CPD 030, Bal-Tec). Glass coverslips were sputter-coated with platinum in a high-vacuum sputtering device (SCD005 sputter coater, Bal-Tec) and imaged with a field emission scanning electron microscope (JSM-6701F,s JEOL) at an acceleration voltage of 8 kV.

### Measurement of RBCs diameter and bleb count on RBCs surface

20 scanning electron micrographs of iRBCs per condition were randomly selected to measure the cell diameter following the scale bar of 1 μm. Additionally, the blebs on the iRBCs surface were counted using ImageJ 1.51k software to determine the drug effect on membrane blebbing.

### Western blot assay

Parasites were synchronised for 2 cycles prior to the start of the experiment. Using 25 cm^2^ flasks, malaria cultures were treated for 10 hours with a final concentration of 1 μM of drug or 1x PBS (control) at 10% parasitemia (mid-late trophozoites) in 2.5% haematocrit. uRBCs cultures were treated only with 1x PBS. Afterward, late stages (schizonts) were enriched using the QuadraMACS Separator and LD columns (Miltenyi Biotec, Germany). A bicinchoninic acid (BCA) assay kit (Thermo Fisher Scientific, USA) was used to determine the final amount of protein per sample. 80 μg of protein were loaded in 8% SDS-PAGE gel and transferred to nitrocellulose membranes. The primary antibodies used for detection were mouse monoclonal anti-μ-calpain (Sigma-Aldrich, United States) diluted 1:1000, mouse monoclonal anti-KAHRP clone 18.2 (European Malaria Reagent Repository, United Kingdom) diluted 1:500 and mouse monoclonal anti-GAPDH (Abcam, United Kingdom) diluted 1:5000. The secondary antibody was goat anti-mouse IgG (H+L) secondary antibody DyLight 680 (LI-COR) diluted 1:5000. Western blot experiments were repeated six times. Image collection was performed using the ChemiDoc MP Imaging System (Bio-Rad, United States). Quantification of KAHRP blots and μ-calpain densitometry plots was done using ImageJ 1.51k software. In order to assess that KAHRP degradation and μ-calpain activation are mediated by calcium redistribution in CQ-, QC- and DCB-treated iRBCs, a pretreatment with the calcium chelator BAPTA-AM (Abcam, United Kingdom) at 50 μM was performed by incubating the parasite cultures at 37 C for 30 min, after which the drugs were added without washing the cells following the protocol described previously.

### EV isolation

Parasites were synchronised for 2 cycles prior to the start of the experiment. Using 75 cm^2^ flasks, malaria cultures were treated for 10 hours with a final concentration of 1 μM of drug or 1x PBS (control) at 10% parasitemia (mid-late trophozoites) in 2.5% haematocrit. uRBCs cultures were treated only with 1x PBS. Briefly, 100 mL of culture growth medium was harvested and centrifuged using the Eppendorf centrifuge 5810 R (Eppendorf, Germany) to remove any additional pellet containing uninfected and iRBCs at 1,800 × *g* for 15 min, twice. This step was followed by centrifugation at 10,000 × *g* for 20 min to remove cell debris using the Allegra X-30R centrifuge (Beckman Coulter, USA). The supernatant was transferred to ultra-clear thin wall tubes (Beckman Coulter, United States) and ultracentrifuged at 200,000 × *g* for 4 hours using a SW 55 Ti rotor in a Beckman OPTIMA90X ultracentrifuge (Beckman Coulter, USA). Finally, 2 washing steps using 1x PBS were performed with the previous ultracentrifugation conditions. The pellet was resuspended in 100 μL 1x PBS and stored at 4C for a maximum of 3 days for characterization and cell-based assays.

### Nanoparticle tracking analysis

The size distribution and concentration of EVs were analyzed using the NanoSight NS300 (Malvern Instruments). Brownian motion of the particles was observed by tracking the scatter signal of individual particles under a microscope. Particle count and number size distribution were obtained. NanoSight measurements gave as a result the particle size and concentration (particle/mL).

### Transmission electron microscopy

The pellet obtained by ultracentrifugation was resuspended with 100 μL of sterile 1x PBS. Samples were fixed with 2.5% glutaraldehyde for 1 hour at 4°C. A volume of 20 μL of each sample was incubated onto a Formvar Film 200 mesh, CU, FF200-Cu grid for 30 min. Negative staining was performed with 5% gadolinium triacetate for 1 min. Samples were viewed using FEI TECNAI SPIRIT G2 RT transmission electron microscopy. (FEI company) across 35,000× magnification.

### Mass spectrometry data acquisition

#### LC-MS

Following the EV isolation protocol described above, after the last washing step, EVs were resuspended in a lysis buffer containing RIPA and protein inhibitor and mixed thoroughly. The samples were incubated for 20 min at 4C, vortex for 15 sec, and stored at −80C until used. EV lysate samples were processed using the S-Trap micro column (Protifi, USA) according to the manufacturer’s recommendations. Total peptide quantification was performed using the Pierce Quantitative Colorimetric Peptide Assay (Thermo Fisher Scientific, USA) to normalise sample peptide concentrations for LCMS analysis. Online reversed-phase (RP) separation of the reconstituted peptides was analysed on the Eksigent Ekspert NanoLC 425 mass spectrometer. Solvent A for RP was 2% acetonitrile, 0.1% formic acid, while solvent B was 98% acetonitrile, 0.1% formic acid. The peptides were first trapped on a Trajan ProteoCol C18P precolumn (3 μm 120 Å, 300 μm × 10 mm) and then resolved on an Acclaim PepMap 100 C18 analytical column (3 μm 100 Å, 75 μm × 250 mm). Peptide elution was performed using a two-step linear gradient, comprising 5 - 12% solvent B over 20 min, followed by 12 - 30% solvent B over 40 min at a flow rate of 300 nL/min. Eluted peptides were directly injected into the TripleTOF 6600 system (SCIEX, USA) for MS-analysis using the information-dependent acquisition (IDA) mode (IonSpray voltage floating 2300 V, interface heater temperature 150 C). Precursor ions were selected across an ion mass of 350 - 1250 m/z with 250 msec accumulation time per spectrum. Maximum 20 precursors were selected for fragmentation, charge state 2-4, intensity > 125 cps with dynamic exclusion for 12 sec after first occurrence and rolling collision energy. MS/MS analysis was performed in high sensitivity mode at 100 msec accumulation time across a mass range of 100 - 1800 m/z.

#### SWATH-MS

For SWATH data acquisition, online RP analyses were done as described for LC-MS. The eluted peptides were injected into a TripleTOF 6600 system (SCIEX, USA) operating in SWATH-MS mode (IonSpray voltage floating 2200 V, interface heater temperature 150 C). Precursor ions were acquired across an ion mass of 400 - 1600 m/z with 50 msec accumulation time per spectrum. Variable window widths were used, specifying a maximum of 100 variable windows across a precursor mass range of 400 - 1200 m/z, with 1 Da window overlap and minimum window width 4 Da. Rolling collision energy was enabled for each window with 5 eV spread. Fragment ion spectra were accumulated in high sensitivity mode for 30 msec over 100 - 1800 m/z mass range.

### Invasion assay with malaria EVs

uRBCs were resuspended to 1% haematocrit with PBS and stained with CellTracker Deep Red (Invitrogen, USA) for 30 mins at RT. To stop the staining reaction, FBS was added followed by 2 washing steps with MCM. A master mix was prepared with stained uRBCs, MCM, and magnet-purified schizont-iRBCs. The master mix was equally distributed in a 96-well flat bottom plate considering duplicates per each condition. EVs from non-treated uRBCs, EVs from non-treated iRBCs and EVs from CQ-treated iRBCs were added containing 300 μg of protein which was calculated using a BCA assay kit (Thermo Fisher Scientific, USA). 1x PBS was used as the control without EVs. For each well, parasites reached 1% parasitemia in 2% haematocrit. The plate was incubated under a gas mixture of 5% CO2, 3% O2, and 92% N2 at 37 °C in the dark for 32 hours. The initial parasitemia was calculated by light microscopy of Giemsa-stained thin smears and flow cytometry. Right after the addition of the EVs (time 0 hours), aliquots of each condition were mixed with PBS and stained with 8 μM Hoechst for 10 min. A washing step was performed with 1x PBS, finally, samples were resuspended with 1x PBS and immediately acquired in a BD LSR Fortessa X-20 (BD, USA). Hoechst staining and flow cytometry acquisition was repeated at 32 hours.

### Stimulation of THP-1-derived macrophages with malaria EVs

The human monocytic leukaemia cell line THP-1 was cultured at 0.5 - 1 × 10^6^ cells/mL in RPMI 1640 Medium, GlutaMAX^™^ Supplement with 10% fetal bovine serum (FBS) (Gibco, Thermo Fisher Scientific, USA) in 25 cm^2^ vented flasks in a humidified incubator at 37°C and 5% CO_2_. THP-1 monocytic cells were transferred to a 12-well flat-bottom plate at 3 × 10^5^ cells/mL. Cells were differentiated using 25 ng/mL phorbol-12-myristate-13-acetate (PMA, Sigma-Aldrich, USA) for 48 hours to obtain a macrophage-like phenotype. The cells were then washed with fresh culture medium and EVs were added at a final concentration of 300 μg of total protein followed by an incubation of 14 hours.

### THP-1-derived macrophages cytokine ELISA assays

IL-1β, TNF-α and IL-6 were measured in culture supernatants from THP-1-derived macrophages upon stimulation with EVs or LPS after 14 hours. Cell culture medium was used as the negative control. These evaluations were performed by the commercial Human ELISA kits (Sigma-Aldrich, USA) according to manufacturer instructions. Experiments were done independently in duplicates. Absorbance was measured by the Infinite^®^ M200 PRO microplate reader (Tecan, Switzerland).

### THP-1-derived macrophages RNA isolation

The medium of the differentiated cells upon stimulation with EVs was carefully removed and kept for cytokine assays. RNA isolation was performed in a sterile area exclusive for RNA work using nuclease free water and RNase-free glass- and plastic ware. Total RNA was isolated by RNAzol (Sigma-Aldrich, USA) following the manufacturer’s protocol. Differentiated cells were mixed with 500 μL of RNAzol to form a homogenous lysate and transferred to 1.5mL tubes. Immediately, 200 μL of RNase-free water was added into the tubes and mixed well followed by an incubation of 10 min at RT. The samples were centrifuged at 12,000 × g for 15 min at RT to separate the semisolid pellet (containing DNA, proteins, and polysaccharides) from the supernatant (containing RNA). 550 μL of supernatant was transferred into new tubes and mixed with equal volume of isopropanol kept at RT for 10 min. Tubes were centrifuged at 12,000× g for 10 min at RT to precipitate RNA. RNA pellet was washed twice with 75% ethanol by centrifugation at 6,000 x g for 2 min at RT. The RNA pellet was air-dried and dissolved in 30 μL of nuclease-free water and stored at −80 C until used.

### THP-1-derived macrophages RNA sequencing

The poly(A) mRNA isolation was performed using Oligo(dT) beads. The mRNA fragmentation was performed using divalent cations and high temperature. Priming was performed using Random Primers. First-strand cDNA and the second-strand cDNA were synthesized. The purified double-stranded cDNA was then treated to repair both ends and add a dA-tailing in one reaction, followed by a T-A ligation to add adaptors to both ends. Size selection of Adaptor-ligated DNA was then performed using DNA Clean Beads. Each sample was then amplified by PCR using P5 and P7 primers and the PCR products were validated. Final libraries with different indexes were multiplexed and loaded on a Novaseq PE150 (Illumina, USA) instrument for sequencing using a 2×150 paired-end (PE) configuration according to manufacturer’s instructions.

### Bioinformatic analysis

#### LC-MS

Acquired spectra were searched using ProteinPilot v5.0, Paragon algorithm v5.0.0.0, 4767 against SwissProt Human reference proteome (UP000005640, 2020 Jun release, 20370 entries) and UniProt Plasmodium falciparum (isolate 3D7) reference proteome (UP000001450, 2020 Jun release, 5456 entries), spiked with common contaminant proteins (cRAP). Cysteine alkyation: methyl methane thiosulfonate (MMTS) was applied and biological modifications were enabled. The Venn diagrams were created in https://bioinformatics.psb.ugent.be/webtools/Venn/ to determine the unique and common proteins between samples. KEGG pathways Over-representation analysis were performed using clusterProfiler (v4.4.4) with significant genes classified as adjusted p-value ≤0.05 with a minimum of 1.5 fold change.

#### SWATH-MS

Acquired spectra were searched using the Spectronaut 15.4.210913.50606 (Biognosys AG, Switzerland), DirectDIA workflow against SwissProt Human reference proteome (UP000005640, 2021 Jan release, 20380 entries) and UniProt *Plasmodium falciparum* (isolate 3D7) reference proteome (UP000001450, 2020 Aug release, 5384 entries), spiked with common contaminant proteins (cRAP). The following modifications were used: MMTS at cysteine as a fixed modification, variable oxidation of methionine and acetylation of the protein N-terminus. The enzyme was set as Trypsin, and up to 2 missed cleavages were permitted. The false discovery rate for peptide and protein identifications was set to 1% and at least 95% confidence, the rest of the parameters were set at the default settings. Briefly, the top 10 peptides based on precursor intensity and confidence were selected across the entire retention time elution range and then scored. The global normalization strategy was based on the median scores. Human and parasite proteome responses were visualized as heatmaps and volcano plots using R package. The latest version of Gene Ontology (GO) annotations for cellular component, biological process and molecular function were obtained as gmt files from gprofiler website (https://biit.cs.ut.ee/gprofiler/gost). The KEGG pathway annotations were mapped directly from KEGG website (https://www.genome.jp/kegg/). Gene Set Enrichment Analysis (GSEA) was performed with R package clusterProfiler (v4.4.4) and the results were visualised with ggplot2. Protein-protein interaction networks were created by submitting the UniProt ascensions to the STRING (Search Tool for the Retrieval of Interacting Genes) software (http://string-db.org/). Interaction networks were visualised for proteins with medium confidence (0.4) with network edges based on evidence, with continuous lines for direct interactions. Clustering was based on an MCL algorithm inflation default parameter of 3.

#### RNA sequencing - THP-1-derived macrophages

In order to remove technical sequences and quality of bases lower than 20, pass filter data were processed by Cutadapt (v1.9.1) to be high-quality clean data. Afterward, the data was aligned to the reference genome via the software Hisat2 (v2.0.1). For the expression analysis, HTSeq (v0.6.1) was used to estimate gene and isoform expression levels from the pair-end clean data. The differential expression analysis was performed by DESeq2 (v1.36.0) Bioconductor package, padj of genes were set <0.05 to detect differential expressed ones. Gene Set Enrichment Analysis (GSEA) was performed with R package clusterProfiler (v4.4.4). Volcano plots with pathway annotations were made with ggplot2 (v3.3.6) and shiny(v1.7.2).

### Statistical analysis

Statistical analysis was performed using GraphPad Prism 9 software (GraphPad Software, USA). Kruskal-Wallis test and two-way analysis of variance (ANOVA) were used to evaluate experiments involving multiple groups. The test used is indicated in the figure legends. Graphs show mean ± SEM. * p < 0.05, ** p < 0.01, *** p < 0.001, **** p < 0.0001.

## DECLARATION OF COMPETING INTEREST

The authors declare that they have no conflicting and/or competing interest in any relevant affiliations or financial involvement with any organization or entity with a financial interest in the subject matter or materials discussed in this manuscript.

## ACKNOWLEDGMENTS

The authors acknowledge the administrative and technical assistance rendered by Ms. Geok Choo Ng at the Department of Microbiology & Immunology, National University of Singapore. The work was funded by generous Tier 1 (R-171-000-098-114) and Tier 3 (R-171-000-143- 592) grants from the Ministry of Education CC-B was supported by the Singapore International Graduate Award (SINGA) Scholarship from A*STAR. Fig. 8 was created with BioRender.com.

## Notes

### Competing Interest Statement

The authors have declared no competing interest.

