## Supplemental Figures for "Chloroquine induces eryptosis in *P. falciparum-infected* red blood cells and the release of extracellular vesicles with a unique protein profile"

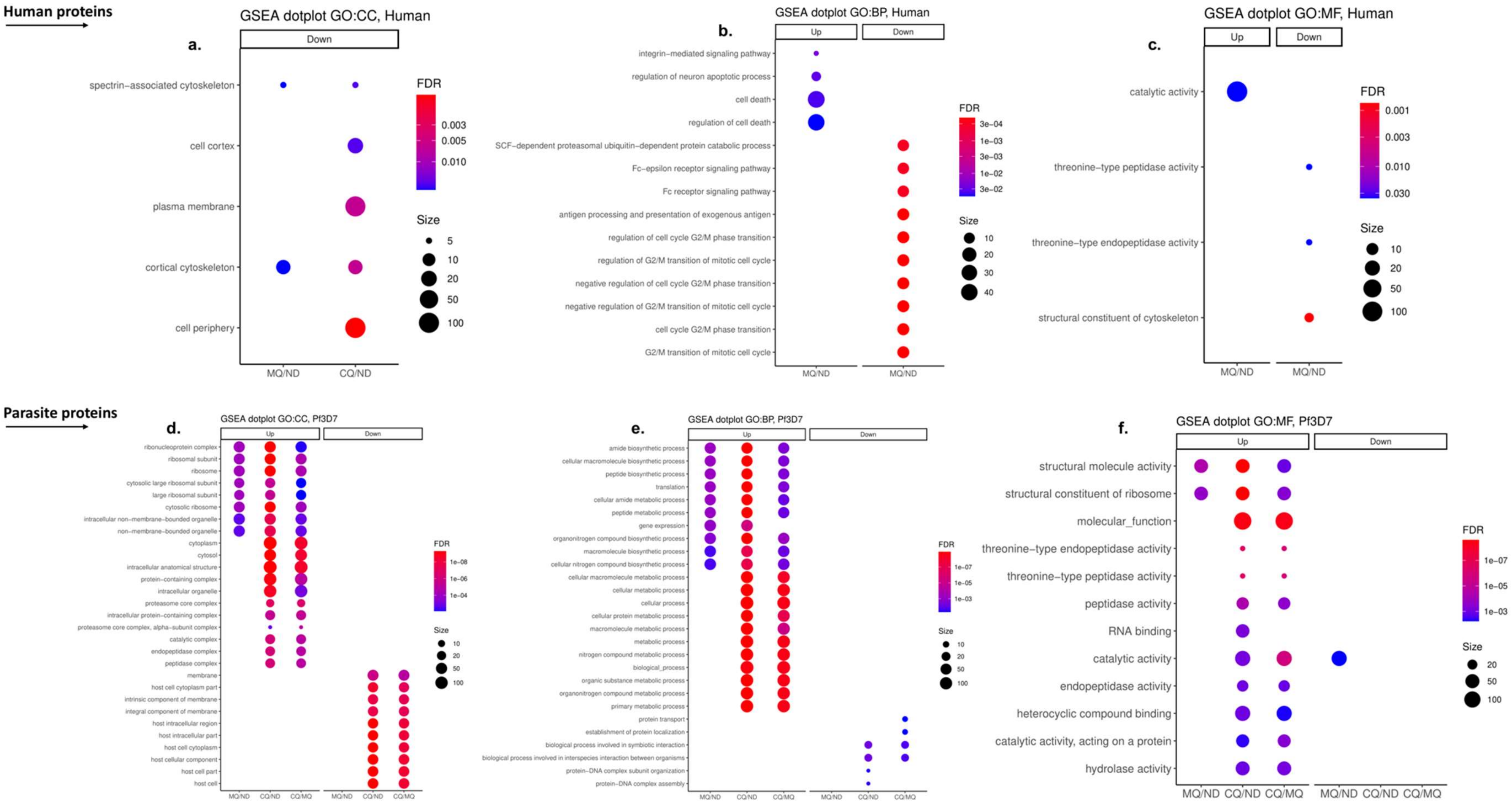


**Supplemental Fig. 1. GO analysis of human and parasite proteins from EVs.** (a & d) Cellular component, (b & e) biological process and (c &f) molecular function. The results represent 3 biological replicates per sample. Only pathways that passed the threshold (fold change ≥ 1.5, adjusted p-value ≤ 0.05) are shown in the bubble plots. ND: EVs from non-treated iRBCs; CQ: EVs from CQ-treated iRBCs; MQ: EVs from MQ-treated iRBCs.

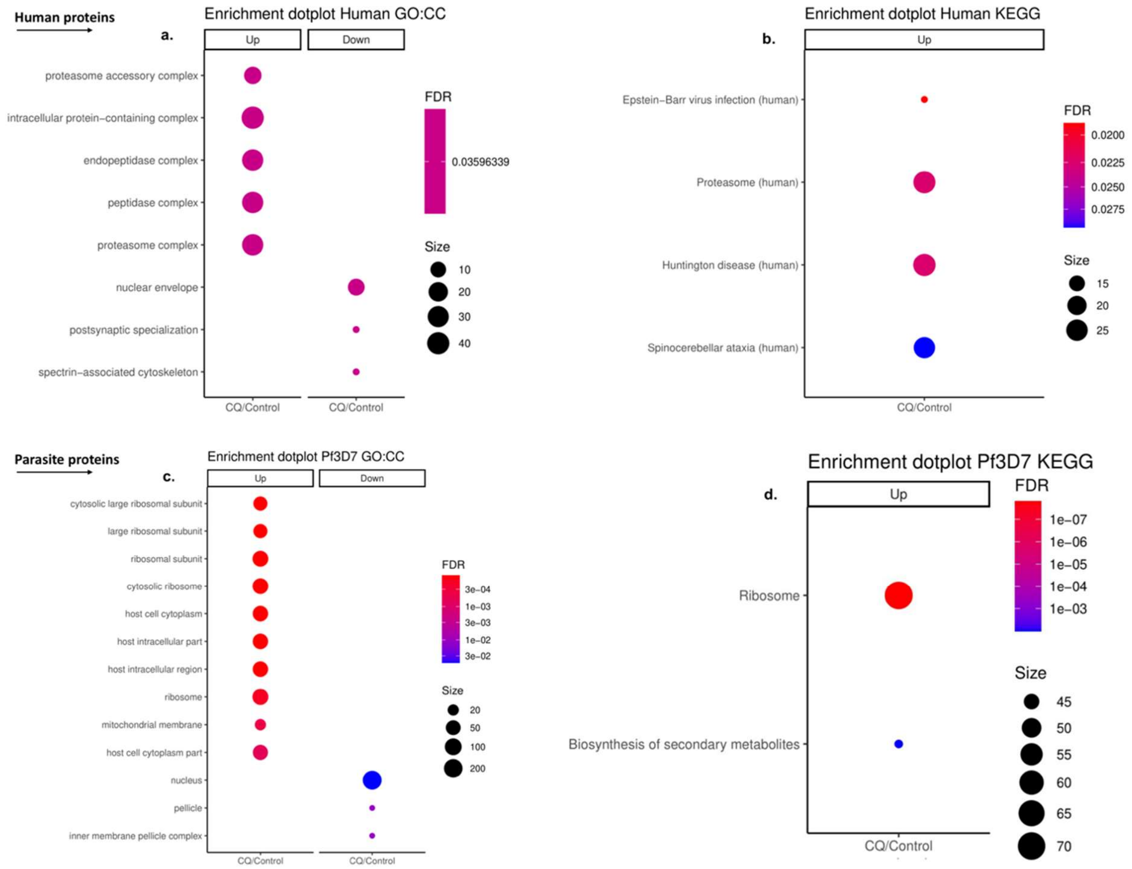


**Supplemental Fig. 2. GO and KEGG analysis of CQ-treated iRBCs lysates.** Human proteins from CQ-treated iRBCs (CQ) in contrast with non-treated iRBCs (Control) were analysed for (a) GO-Cellular component and (b) KEGG enrichment pathways. Parasite proteins from CQ-treated iRBCs in contrast with non-treated iRBCs were analysed for (c) GO-Cellular component and (d) KEGG enrichment pathways. The results represent 3 biological replicates per sample. Only pathways that passed the threshold (fold change ≥ 1.5, adjusted p-value ≤ 0.05) are shown in the bubble plots.

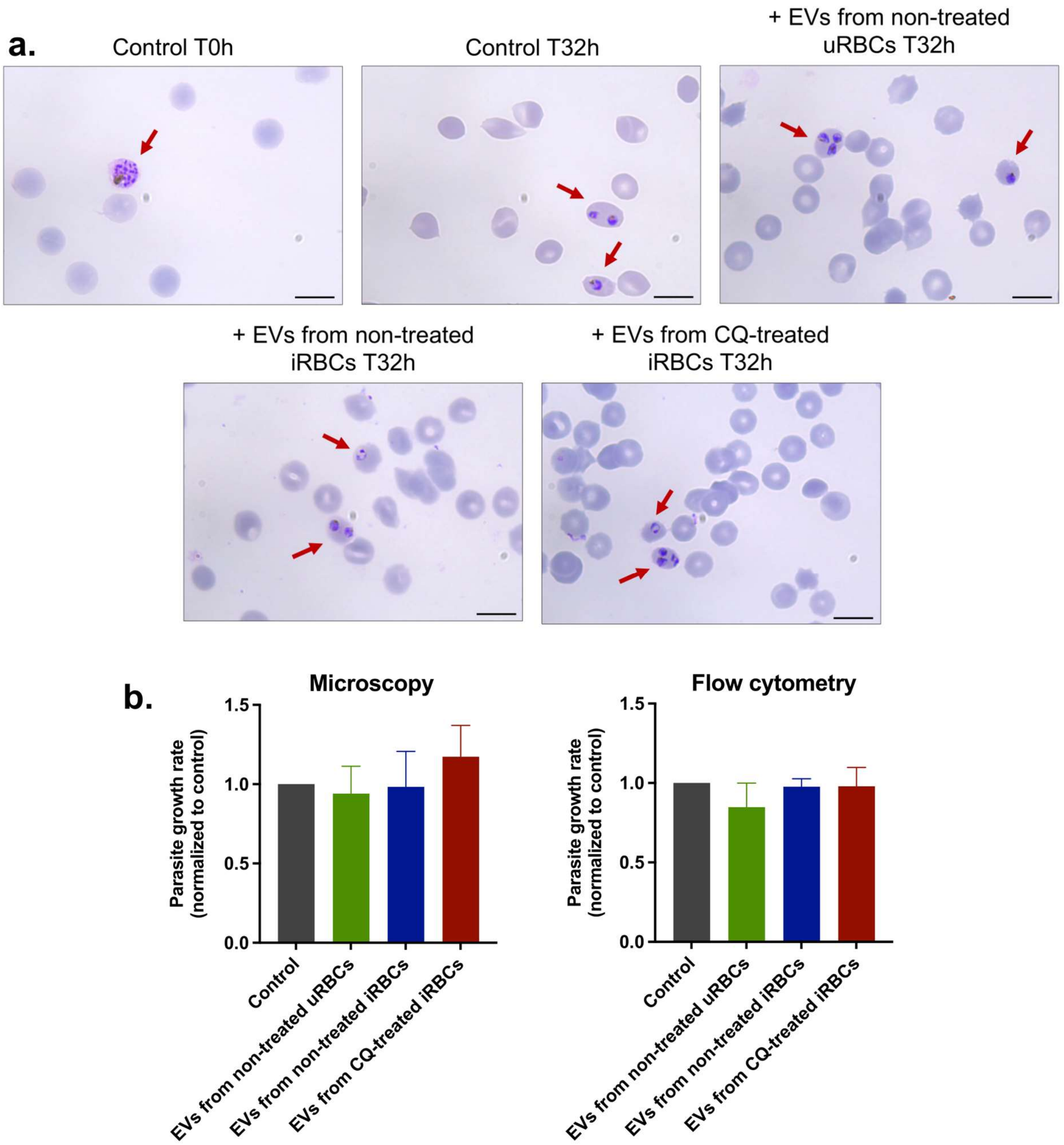


**Supplemental Fig. 3. iRBCs-derived EVs effect in parasite growth rate.** (a) Giemsa smears of invasion assays at T0h and T32h. 1X PBS used to resuspend EV samples was included as the control. Scale bars = 10 μm. (c) Parasite growth rate normalized to the control from microscopy and flow cytometry data. The results are represented in mean ± s.e.m. of 3 independent experiments by Kruskal-Wallis test with post-hoc Dunn's test. No significant difference was found.

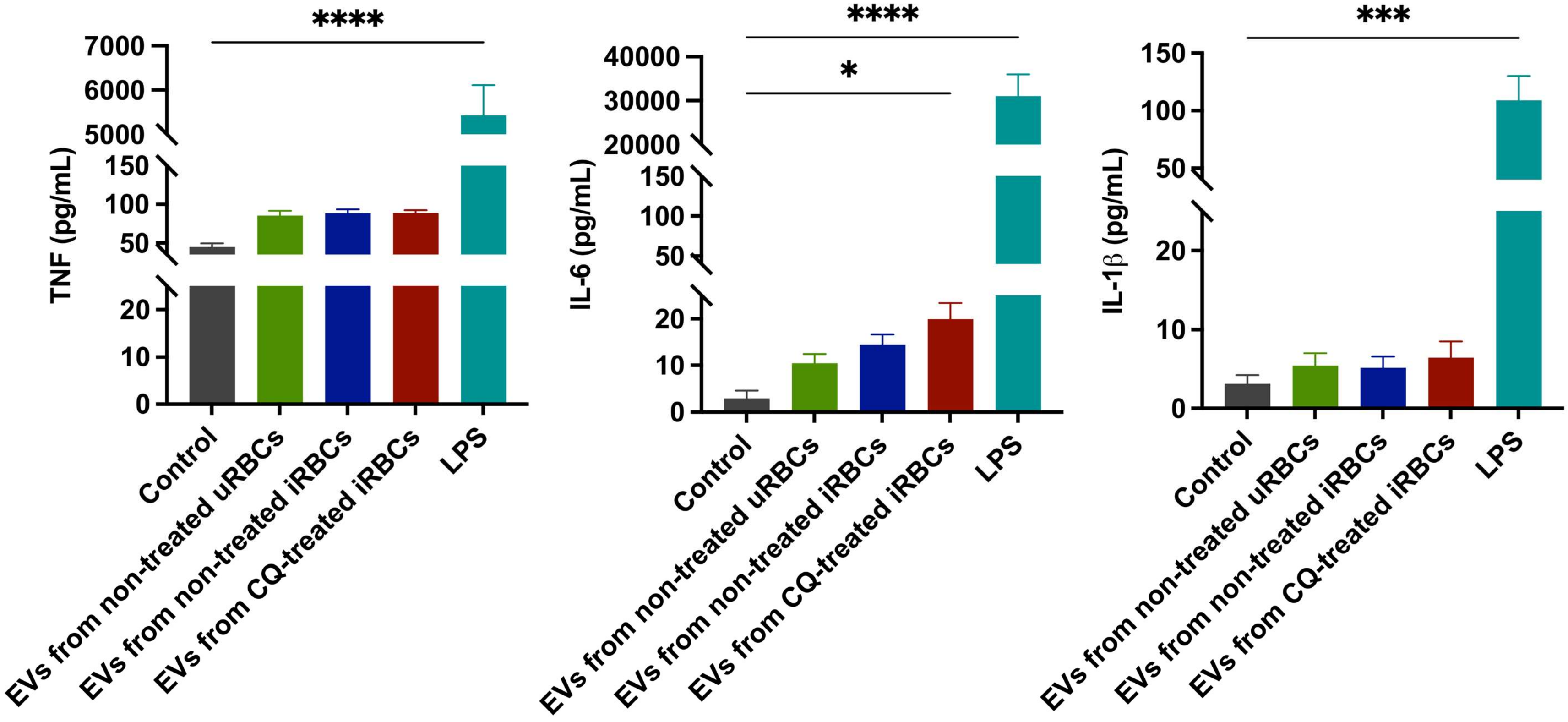


**Supplemental Fig. 4. ELISA assay for detection of TNF, IL-6 and IL-1β in THP-1 derived-macrophages.** All samples were compared to the control (media without EVs). LPS was used as a positive control. The results are represented in mean ± SEM of 3 independent experiments; *$p < 0.05$, ****$p < 0.0001$ by Kruskal-Wallis test with post-hoc Dunn's test.

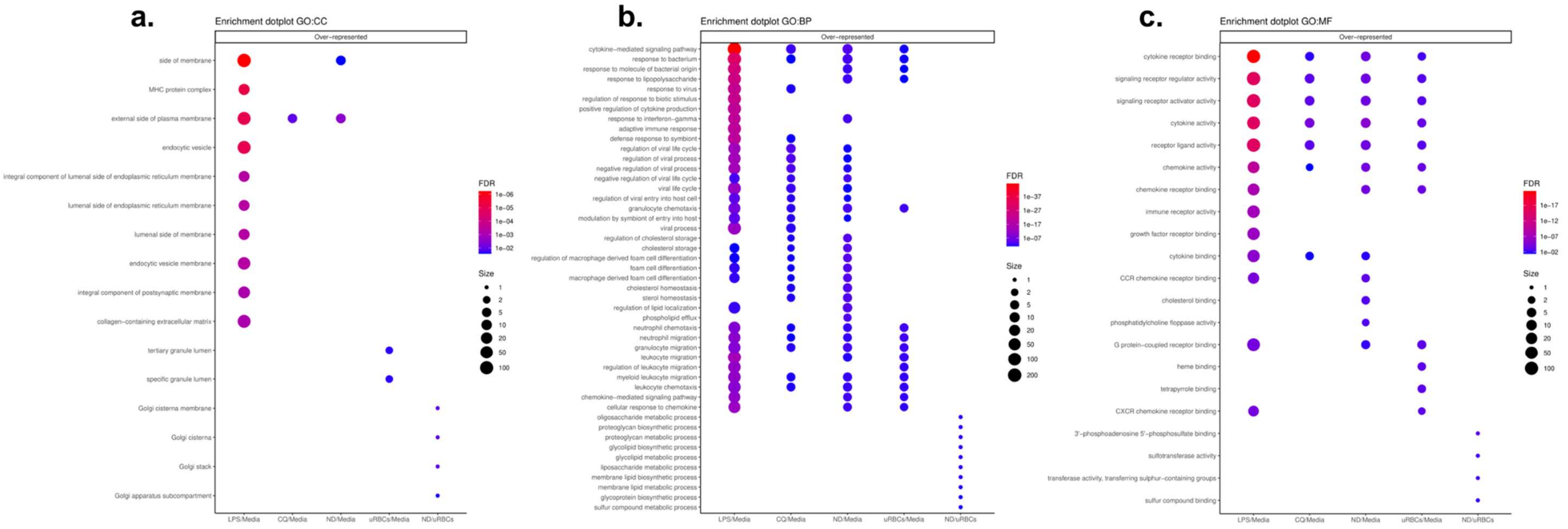


**Supplemental Fig. 5. GO analysis of THP-1-derived macrophages upon stimulation with EVs.** (a) Cellular component, (b) biological process and (c) molecular function. The results represent 3 biological replicates per sample. Only pathways that passed the threshold (fold change ≥ 1.5, adjusted p-value ≤ 0.05) are shown in the bubble plots
