## Supplemental Table 1 for "Chloroquine induces eryptosis in *P. falciparum-infected* red blood cells and the release of extracellular vesicles with a unique protein profile"

**Pathway enrichment analysis of parasite and human SWATH-MS datasets**

| Description | Contrast | Test | Organism | Sign | Set size | Enrichment score | NES | pvalue | p.adjust | qvalues | Rank | Core_enrichment |
|---|---|---|---|---|---|---|---|---|---|---|---|---|
| Proteasome core complex | CQ/ND | GO:CC | Pf3D7 | Up | 14 | 0.843668054 | 2.662333865 | 2.48E-08 | 8.04E-08 | 2.24E-08 | 40 | PF3D7_1328100/PF3D7_1011400/PF3D7_0317000/PF3D7_0727400/PF3D7_0807500/PF3D7_1353800/PF3D7_0931800/PF3D7_1470900/PF3D7_1353900/PF3D7_0518300/PF3D7_0608500/PF3D7_0803800/PF3D7_1474800 |
| | CQ/MQ | GO:CC | Pf3D7 | Up | 14 | 0.852541461 | 2.495291371 | 3.88E-08 | 2.03E-07 | 7.55E-08 | 21 | PF3D7_1328100/PF3D7_1353800/PF3D7_0518300/PF3D7_1011400/PF3D7_0608500/PF3D7_0317000/PF3D7_0807500/PF3D7_0803800/PF3D7_1353900/PF3D7_0727400/PF3D7_0931800/PF3D7_1470900/PF3D7_1474800 |
| | MQ/ND | GO:CC | Pf3D7 | Down | 14 | -0.401707203 | -1.010309205 | 0.462382445 | 0.628840125 | 0.506187098 | 68 | PF3D7_1470900/PF3D7_0727400/PF3D7_1011400/PF3D7_0931800/PF3D7_1353900/PF3D7_0807500/PF3D7_1474800/PF3D7_0803800/PF3D7_0317000/PF3D7_0608500/PF3D7_0518300/PF3D7_1353800/PF3D7_1328100 |
| Proteasome core complex, alpha-subunit complex | CQ/ND | GO:CC | Pf3D7 | Up | 7 | 0.816666667 | 2.069180917 | 0.000761535 | 0.001232961 | 0.00034355 | 40 | PF3D7_0317000/PF3D7_0727400/PF3D7_0807500/PF3D7_1353800/PF3D7_1353900/PF3D7_0608500/PF3D7_1474800 |
| | CQ/MQ | GO:CC | Pf3D7 | Up | 7 | 0.922222222 | 2.184279729 | 1.43E-06 | 6.49E-06 | 2.41E-06 | 21 | PF3D7_1353800/PF3D7_0608500/PF3D7_0317000/PF3D7_0807500/PF3D7_1353900/PF3D7_0727400/PF3D7_1474800 |
| | MQ/ND | GO:CC | Pf3D7 | Down | 7 | -0.694444444 | -1.46616737 | 0.057660626 | 0.204233096 | 0.164398157 | 63 | PF3D7_1353900/PF3D7_0807500/PF3D7_1474800/PF3D7_0317000/PF3D7_0608500/PF3D7_1353800 |
| Proteasome | CQ/ND | KEGG | Pf3D7 | Up | 25 | 0.706694486 | 2.589158028 | 1.85E-07 | 6.46E-07 | 4.86E-07 | 29 | PF3D7_1328100/PF3D7_1011400/PF3D7_0317000/PF3D7_0727400/PF3D7_0807500/PF3D7_1353800/PF3D7_0931800/PF3D7_1470900/PF3D7_1353900/PF3D7_0518300/PF3D7_0608500/PF3D7_0803800 |
| | CQ/MQ | KEGG | Pf3D7 | Up | 25 | 0.691196774 | 2.344698735 | 2.37E-06 | 1.66E-05 | 2.50E-06 | 21 | PF3D7_1328100/PF3D7_1353800/PF3D7_0518300/PF3D7_1011400/PF3D7_0608500/PF3D7_0317000/PF3D7_0807500/PF3D7_0803800/PF3D7_1353900/PF3D7_0727400/PF3D7_0931800/PF3D7_1470900/PF3D7_1474800 |
| | MQ/ND | KEGG | Pf3D7 | Down | 25 | -0.485939649 | -1.438170266 | 0.063400576 | 0.147934678 | 0.111229081 | 74 | PF3D7_0413600/PF3D7_0205900/PF3D7_1470900/PF3D7_1248900/PF3D7_1130400/PF3D7_0727400/PF3D7_1011400/PF3D7_0931800/PF3D7_1368100/PF3D7_1353900/PF3D7_1306400/PF3D7_0807500/PF3D7_1402300/PF3D7_1338100/PF3D7_1466300/PF3D7_1474800/PF3D7_0803800/PF3D7_0317000/PF3D7_1311500/PF3D7_0608500/PF3D7_0518300/PF3D7_1353800/PF3D7_1328100 |
| Ribosome | CQ/ND | KEGG | Pf3D7 | Up | 45 | 0.716531657 | 2.960852635 | 1.00E-10 | 7.00E-10 | 5.26E-10 | 78 | PF3D7_1109900/PF3D7_1323400/PF3D7_1027800/PF3D7_1358800/PF3D7_1126200/PF3D7_1302800/PF3D7_0618300/PF3D7_0309600/PF3D7_1317800/PF3D7_0317600/PF3D7_1408600/PF3D7_1465900/PF3D7_0520000/PF3D7_1105400/PF3D7_1341200/PF3D7_0516900/PF3D7_0813900/PF3D7_1004000/PF3D7_0422400/PF3D7_1460700/PF3D7_1424100/PF3D7_1142600/PF3D7_0415900/PF3D7_1447000/PF3D7_0719600/PF3D7_1309100/PF3D7_1431700/PF3D7_0322900/PF3D7_1426000/PF3D7_1351400/PF3D7_1338200/PF3D7_0507100/PF3D7_1130200/PF3D7_0519400/PF3D7_1414300/PF3D7_1019400/PF3D7_0821700/PF3D7_1331800/PF3D7_1323100/PF3D7_1342000/PF3D7_0307200/PF3D7_1341300 |
| | CQ/MQ | KEGG | Pf3D7 | Up | 45 | 0.554569835 | 2.113614655 | 2.80E-05 | 9.78E-05 | 1.47E-05 | 101 | PF3D7_1331800/PF3D7_1126200/PF3D7_1317800/PF3D7_1130200/PF3D7_1358800/PF3D7_0519400/PF3D7_0422400/PF3D7_1302800/PF3D7_0309600/PF3D7_1465900/PF3D7_0520000/PF3D7_0317600/PF3D7_1408600/PF3D7_0813900/PF3D7_1105400/PF3D7_1323400/PF3D7_1447000/PF3D7_0719600/PF3D7_1027800/PF3D7_0322900/PF3D7_1424100/PF3D7_0618300/PF3D7_1341300/PF3D7_1460700/PF3D7_0821700/PF3D7_1019400/PF3D7_1341200/PF3D7_0516900/PF3D7_1004000/PF3D7_1431700/PF3D7_1309100/PF3D7_0415900/PF3D7_1426000/PF3D7_1323100/PF3D7_1338200/PF3D7_1109900/PF3D7_1142600/PF3D7_1351400/PF3D7_1414300/PF3D7_0507100 |
| | MQ/ND | KEGG | Pf3D7 | Up | 45 | 0.570951849 | 2.244802489 | 4.67E-06 | 3.27E-05 | 2.46E-05 | 36 | PF3D7_1109900/PF3D7_1027800/PF3D7_1342000/PF3D7_1323400/PF3D7_1142600/PF3D7_0618300/PF3D7_0507100/PF3D7_1341200/PF3D7_1004000/PF3D7_0516900/PF3D7_1351400/PF3D7_0614500/PF3D7_0415900/PF3D7_1338200/PF3D7_1414300/PF3D7_0307200/PF3D7_1426000/PF3D7_1309100/PF3D7_1431700/PF3D7_1424400 |
| Proteasome (human) | CQ/ND | KEGG | Human | Up | 21 | 0.501670234 | 1.349004967 | 0.118343195 | 0.746872587 | 0.746872587 | 101 | PSMC6/PSMC2/PSME2/PSMC3/PSMD3/PSMD11/PSMB4/PSMB1/PSMC5/PSMD2/PSMA6/PSMB3/PSMB2/PSMA5/PSMA4/PSMB5/PSMA2/PSMA3 |
| | CQ/MQ | KEGG | Human | Up | 21 | 0.69034724 | 1.814539442 | 0.00151531 | 0.039398056 | 0.036154761 | 63 | PSME2/PSMB4/PSMB1/PSMA2/PSMD2/PSMB3/PSMB5/PSMC6/PSMA6/PSMA4/PSMB2/PSMA5/PSMA3/PSMD12/PSMC2/PSMD3/PSMC5/PSMA1/PSMC3 |
| | MQ/ND | KEGG | Human | Down | 21 | -0.304372506 | -1.19405531 | 0.134927247 | 0.473578595 | 0.409028746 | 43 | PSMA1/PSMD12/PSMA3/PSMA5/PSMB2/PSMA6/PSMA4/PSME2/PSMB4/PSMD2/PSMB1/PSMB3/PSMB5/PSMA2 |

Upregulated

Downregulated
