## Supplemental Table 2 for "Chloroquine induces eryptosis in *P. falciparum-infected* red blood cells and the release of extracellular vesicles with a unique protein profile"

**Differentially expressed human proteasome subunits in EV samples.**

| Gene ID | Protein name - Human | CQ/ND | | CQ/MQ | | MQ/ND | |
|---|---|---|---|---|---|---|---|
| | | adjusted p-value | log2(FC) | adjusted p-value | log2(FC) | adjusted p-value | log2(FC) |
| PSMD11 | 26S proteasome non-ATPase regulatory subunit 11 | 0.0183 | 0.7244 | 0.0687 | 0.2197 | 0.0451 | 0.5047 |
| PSMD2 | 26S proteasome non-ATPase regulatory subunit 2 | 0.0246 | 0.6085 | 0.0069 | 0.8420 | 0.1639 | -0.2335 |
| PSMC3 | 26S proteasome regulatory subunit 6A | 0.0176 | 0.7606 | 0.0787 | 0.3188 | 0.0573 | 0.4419 |
| PSMC2 | 26S proteasome regulatory subunit 7 | 0.0240 | 0.8101 | 0.0718 | 0.4787 | 0.0568 | 0.3314 |
| PSMC5 | 26S proteasome regulatory subunit 8 | 0.0161 | 0.6716 | 0.0745 | 0.4019 | 0.0406 | 0.2697 |
| PSME2 | Proteasome activator complex subunit 2 | 0.0397 | 0.7733 | 0.0084 | 0.9921 | 0.0164 | -0.2188 |
| PSMF1 | Proteasome inhibitor PI31 subunit | 0.1373 | 0.2729 | 0.0073 | -1.0523 | 0.0523 | 1.3252 |
| PSMA2 | Proteasome subunit alpha type-2 | 0.1089 | 0.4651 | 0.0449 | 0.8821 | 0.0978 | -0.4170 |
| PSMB1 | Proteasome subunit beta type-1 | 0.0942 | 0.6722 | 0.0465 | 0.9092 | 0.1018 | -0.2370 |
| PSMB2 | Proteasome subunit beta type-2 | 0.0877 | 0.5413 | 0.0433 | 0.7154 | 0.1533 | -0.1741 |
| PSMB4 | Proteasome subunit beta type-4 | 0.0778 | 0.7058 | 0.0399 | 0.9366 | 0.1566 | -0.2308 |

Upregulated

Downregulated
