## Supplemental Table 3 for "Chloroquine induces eryptosis in *P. falciparum-infected* red blood cells and the release of extracellular vesicles with a unique protein profile"

**Differentially expressed parasite ribosomal proteins and proteasome subunits in EV samples.**

| Gene ID | Protein name - Human | CQ/ND | | CQ/MQ | | MQ/ND | |
|---|---|---|---|---|---|---|---|
| | | adjusted p-value | log2(FC) | adjusted p-value | log2(FC) | adjusted p-value | log2(FC) |
| PF3D7_0317600 | 40S ribosomal protein S11, putative | 0.0249 | 1.0364 | 0.0277 | 0.9968 | 0.2489 | 0.0395 |
| PF3D7_1358800 | 40S ribosomal protein S15 | 0.0029 | 1.3799 | 0.0071 | 1.2157 | 0.1018 | 0.1642 |
| PF3D7_0813900 | 40S ribosomal protein S16, putative | 0.0299 | 0.9147 | 0.0377 | 0.8501 | 0.2217 | 0.0645 |
| PF3D7_1126200 | 40S ribosomal protein S18, putative | 0.0091 | 1.2135 | 0.0048 | 1.3713 | 0.0848 | -0.1578 |
| PF3D7_1317800 | 40S ribosomal protein S19 | 0.0164 | 1.0929 | 0.0063 | 1.3561 | 0.0827 | -0.2632 |
| PF3D7_0422400 | 40S ribosomal protein S19 | 0.2106 | 0.8941 | 0.0263 | 1.1879 | 0.0146 | -0.2937 |
| PF3D7_0519400 | 40S ribosomal protein S24 | 0.0209 | 0.6979 | 0.0008 | 1.2032 | 0.0459 | -0.5053 |
| PF3D7_1465900 | 40S ribosomal protein S3 | 0.0165 | 1.0007 | 0.0142 | 1.0225 | 0.2255 | -0.0218 |
| PF3D7_1105400 | 40S ribosomal protein S4 | 0.0077 | 0.9663 | 0.0135 | 0.8365 | 0.1973 | 0.1299 |
| PF3D7_1302800 | 40S ribosomal protein S7 | 0.0106 | 1.1484 | 0.0090 | 1.1460 | 0.2275 | 0.0024 |
| PF3D7_1408600 | 40S ribosomal protein S8 | 0.0170 | 1.0297 | 0.0280 | 0.9329 | 0.1633 | 0.0968 |
| PF3D7_0520000 | 40S ribosomal protein S9, putative | 0.0255 | 0.9795 | 0.0277 | 1.0061 | 0.2409 | -0.0266 |
| PF3D7_1130200 | 60S acidic ribosomal protein P0 | 0.0642 | 0.7157 | 0.0093 | 1.2912 | 0.0662 | -0.5755 |
| PF3D7_0309600 | 60S acidic ribosomal protein P2 | 0.0070 | 1.0995 | 0.0092 | 1.0898 | 0.1945 | 0.0097 |
| PF3D7_0719600 | 60S ribosomal protein L11a, putative | 0.0427 | 0.7678 | 0.0390 | 0.6963 | 0.2360 | 0.0715 |
| PF3D7_1004000 | 60S ribosomal protein L13, putative | 0.0343 | 0.8950 | 0.1397 | 0.4474 | 0.1274 | 0.4476 |
| PF3D7_1341200 | 60S ribosomal protein L18a | 0.0405 | 0.9500 | 0.1559 | 0.5021 | 0.1258 | 0.4479 |
| PF3D7_1323400 | 60S ribosomal protein L23 | 0.0424 | 1.3939 | 0.2219 | 0.8109 | 0.0422 | 0.5831 |
| PF3D7_1331800 | 60S ribosomal protein L23, putative | 0.0441 | 0.6205 | 0.0026 | 1.4503 | 0.0382 | -0.8298 |
| PF3D7_1460700 | 60S ribosomal protein L27 | 0.0408 | 0.8783 | 0.1001 | 0.5863 | 0.1764 | 0.2920 |
| PF3D7_0618300 | 60S ribosomal protein L27a, putative | 0.0110 | 1.1146 | 0.1327 | 0.6398 | 0.1236 | 0.4747 |
| PF3D7_1027800 | 60S ribosomal protein L3 | 0.0192 | 1.3820 | 0.1196 | 0.6949 | 0.0966 | 0.6871 |
| PF3D7_1142600 | 60S ribosomal protein L35ae, putative | 0.0437 | 0.8091 | 0.1964 | 0.3250 | 0.1353 | 0.4841 |
| PF3D7_1109900 | 60S ribosomal protein L36 | 0.0201 | 1.5816 | 0.2242 | 0.3322 | 0.0608 | 1.2495 |
| PF3D7_0415900 | Ribosomal protein L15 | 0.0386 | 0.8068 | 0.1530 | 0.3998 | 0.1419 | 0.4070 |
| PF3D7_0727400 | Proteasome subunit alpha type | 0.0431 | 1.2130 | 0.0209 | 1.5020 | 0.1125 | -0.2890 |
| PF3D7_1353900 | Proteasome subunit alpha type | 0.0491 | 1.1774 | 0.0229 | 1.5287 | 0.0941 | -0.3513 |
| PF3D7_1353800 | Proteasome subunit alpha type | 0.1045 | 1.1859 | 0.0094 | 1.9544 | 0.0190 | -0.7685 |
| PF3D7_1474800 | Proteasome subunit alpha type-1, putative | 0.0725 | 0.9341 | 0.0255 | 1.4054 | 0.0743 | -0.4713 |
| PF3D7_0608500 | Proteasome subunit alpha type-2, putative | 0.0576 | 1.1090 | 0.0216 | 1.7221 | 0.0618 | -0.6132 |
| PF3D7_0317000 | Proteasome subunit alpha type-3, putative | 0.0484 | 1.2214 | 0.0228 | 1.7027 | 0.0635 | -0.4813 |
| PF3D7_0807500 | Proteasome subunit alpha type-6, putative | 0.0545 | 1.2124 | 0.0268 | 1.5892 | 0.1037 | -0.3768 |
| PF3D7_1011400 | Proteasome subunit beta | 0.0157 | 1.4880 | 0.0092 | 1.7827 | 0.0982 | -0.2948 |
| PF3D7_1470900 | Proteasome subunit beta | 0.0269 | 1.1777 | 0.0119 | 1.4436 | 0.0989 | -0.2660 |
| PF3D7_1328100 | Proteasome subunit beta | 0.0323 | 1.5477 | 0.0129 | 2.3204 | 0.0592 | -0.7726 |
| PF3D7_0518300 | Proteasome subunit beta | 0.0474 | 1.1543 | 0.0194 | 1.7914 | 0.0985 | -0.6370 |
| PF3D7_0803800 | Proteasome subunit beta | 0.0588 | 1.0584 | 0.0209 | 1.5390 | 0.0774 | -0.4806 |
| PF3D7_0108000 | Proteasome subunit beta | 0.1038 | -0.0041 | 0.0302 | -2.4508 | 0.0233 | 2.4467 |
| PF3D7_0931800 | Proteasome subunit beta type-6, putative | 0.0371 | 1.1801 | 0.0197 | 1.4812 | 0.1057 | -0.3011 |

Upregulated

Downregulated
