## Supplemental Table 4 for "Chloroquine induces eryptosis in *P. falciparum-infected* red blood cells and the release of extracellular vesicles with a unique protein profile"

**Differentially expressed genes from KEGG and Reactome pathways of THP-1-derived macrophages upon stimulations with EVs**

| | | uRBCs/Media | | ND/Media | | CQ/Media | |
|---|---|---|---|---|---|---|---|
| **Pathway enrichment analysis** | **Gene ID** | **adjusted p-value** | **log2(FC)** | **adjusted p-value** | **log2(FC)** | **adjusted p-value** | **log2(FC)** |
| KEGG: Cytokine-cytokine receptor interaction (human) | CCL20 | 2.76E-09 | 0.7454 | 4.81E-18 | 0.9990 | 1.80E-11 | 0.8257 |
| | CXCL1 | 4.52E-04 | 0.7437 | 4.58E-07 | 0.9251 | 5.83E-05 | 0.7973 |
| | IL18R1 | 3.83E-02 | 1.0492 | 2.97E-02 | 0.9193 | 7.09E-03 | 1.1485 |
| | CXCL10 | 1.83E-02 | 2.0927 | 5.61E-01 | 0.6531 | 1.58E-01 | 1.3881 |
| | CSF1 | 1.89E-02 | 0.4904 | 3.22E-05 | 0.6685 | 6.25E-06 | 0.7242 |
| | TNFRSF9 | 7.01E-02 | 0.7420 | 1.75E-04 | 1.0661 | 4.86E-03 | 0.9062 |
| | CCL4L2 | 8.48E-02 | 0.7244 | 3.05E-03 | 0.8863 | 3.47E-01 | 0.4416 |
| | EBI3 | 1.13E-01 | 0.7705 | 4.86E-03 | 0.9619 | 8.46E-03 | 0.9840 |
| | IL32 | 4.17E-01 | 0.7089 | 8.56E-03 | 1.1464 | 4.73E-02 | 1.0077 |
| | CCL2 | 6.37E-02 | 1.1791 | 2.44E-02 | 1.1303 | 6.19E-02 | 1.0689 |
| | OSMR | 4.58E-01 | -0.3842 | 1.01E-03 | -0.7868 | 3.37E-02 | -0.6005 |
| | IL11RA | 7.09E-01 | -0.3783 | 2.59E-02 | -0.7514 | 5.82E-02 | -0.7212 |
| Reactome: Interferon alpha/beta signaling | MX1 | 5.74E-02 | 1.0352 | 4.00E-01 | 0.4780 | 3.14E-03 | 1.2693 |
| | IFIT1 | 7.46E-02 | 1.0664 | 3.38E-01 | 0.5723 | 2.35E-02 | 1.1265 |
| | IFITM1 | 6.29E-02 | 1.2428 | NA | 0.7900 | 7.22E-03 | 1.4394 |

Upregulated

Downregulated

NA: Not accurate (low gene expression levels)
